# Identification of a novel mycovirus belonging to the “flexivirus”-related family with icosahedral virion

**DOI:** 10.1101/2024.09.25.614901

**Authors:** Chien-Fu Wu, Ryo Okada, Uri Neri, Yi-Cheng Chang, Ogawara Takashi, Kentaro Kitaura, Ken Komatsu, Hiromitsu Moriyama

**Affiliations:** Laboratory of Molecular and Cellular Biology, Graduate School of Agriculture, Tokyo University of Agriculture and Technology, 3-5-8 Saiwaicho, Fuchu, Tokyo 183-8509, Japan; Horticultural Research Institute, Ibaraki Agricultural Center, 3165-1 Ago, Kasama 319-0292, Japan; The Shmunis School of Biomedicine and Cancer Research, Tel Aviv University, Tel Aviv 6997801, Israel; Laboratory of Plant Pathology, Graduate School of Agriculture, Tokyo University of Agriculture and Technology, 3-5-8 Saiwaicho, Fuchu, Tokyo 183-8509, Japan; Institute of Global Innovation Research (GIR), Tokyo University of Agriculture and Technology (TUAT), 3-5-8 Saiwaicho, Fuchu, Tokyo 183-8509, Japan

**Keywords:** coat protein, single jelly-roll, virus particle, mycovirus, deltaflexivirus, *Tymovirales*

## Abstract

The order *Tymovirales* currently comprises five viral families with positive-sense RNA ((+)RNA) genomes that infect plants, fungi, and insects. Virion morphologies within the order *Tymovirales* differ between families, with icosahedral virions in the *Tymoviridae* and filamentous virions in the other “*flexi*”*viridae* families. Despite their different morphologies, these viruses are placed in the same order based on phylogenetic analyses of replicase-associated polyproteins. However, one of the families in the *Tymovirales, Deltaflexiviridae*, is considered to be capsidless because there have been no published reports of virion isolation. Here we report that a new “flexivirus”-related (+)RNA virus, prospectively named <u>F</u>usarium <u>o</u>xysporum icosahedral <u>v</u>irus <u>1</u> (FoIV1), is icosahedral, and that most deltaflexiviruses may have icosahedral virions. Phylogenetic analyses based on replicase-associated polyproteins indicated that FoIV1 forms a distinct group in the *Tymovirales* with some viruses originally assigned to the *Deltaflexiviridae*. Electron microscopy, protein analysis, and protein structure predictions indicate that FoIV1 ORF4 encodes a single jelly-roll (SJR)-like coat protein (CP) that constitutes the icosahedral virions. Results of clustering analyses based on amino acid sequences and predicted CP structures suggested that most of the deltaflexiviruses have icosahedral virions composed of SJR-like CPs as in FoIV1, rather than having filamentous virions or capsidless. These results challenge the conventional understanding of viruses in the order *Tymovirales*, with important implications for revising its taxonomic framework and providing insights into the evolutionary relationships within this diverse and broad host range group of (+)RNA viruses.

## Introduction

Mycoviruses, also known as fungal viruses, have been discovered in a number of fungal groups, as well as yeasts and oomycetes (water moulds). They generally fall into the double-stranded RNA (dsRNA), single-stranded with either positive or negative-strand RNA ((+/-)RNA), reverse transcribing (RT) RNA viruses, or single-stranded DNA (ssDNA) genomes, with the majority of them being either dsRNA or (+)RNA genomes (Ghabrial et al., 2015). While most mycoviruses are latent, some influence host phenotypes. For example, some mycoviruses can impair growth, alter pigmentation, and sporulation or reduce virulence among pathogenic fungi (Ghabrial et al., 2015; Kotta-Loizou, 2021). These properties make mycoviruses promising candidates for controlling fungal plant diseases, as exemplified by the successful use of hypovirulent *Cryphonectria parasitica* isolates infected with Cryphonectria parasitica hypovirus 1 (CHV1) to manage chestnut blight (Nuss, 2005; Heiniger and Rigling, 2009; Robin et al., 2010; Prospero and Rigling, 2016).

The advent of high-throughput sequencing (HTS) has revolutionized virus discovery, including the identification of numerous novel mycoviruses (Marzano et al., 2016; Marzano and Domier, 2016; Bartholomäus et al., 2016; Ruiz-Padilla et al., 2021; Raco et al., 2022). Concurrently, advances in bioinformatics have facilitated viral diversity analysis, large scale virus identification (Edgar et al., 2022; Neri et al., 2022), protein structure prediction (Jumper et al., 2021), and the discovery of hidden motifs in viral replicases (Urayama et al., 2024). However, certain molecular properties, such as viral RNA structures, are challenging to reliably predict *in silico* and therefore require experimental verification. In the endornaviruses, a group of large (+)RNA viruses (Roossinck et al., 2011), a site-specific nick near the 5’-terminus of the coding strand has been demonstrated by northern hybridization (Fukuhara et al., 1995; Okada et al., 2011; Valverde et al., 2019). Viruses belonging to the family *Alternaviridae*, which was established in 2022 (Kotta-Loizou et al., 2022), are the first group of dsRNA viruses whose genome segments known to be 5’-7-methylguanosine (m^7^G)-capped and 3’-polyadenylated. The presence of 5’-end cap structures was confirmed using RNA dot blots with anti-cap antibodies or the oligo-capping method (Wu et al., 2021; Lutz et al., 2022). Furthermore, although an increasing number of mycovirus genomes have been sequenced, and the functions of most viral-encoded proteins have been predicted based on sequence similarity with known proteins, there are still many open reading frames (ORFs) in the genomes of mycoviruses whose translation products (if any) have not yet been investigated, often having low or no sequence similarity to proteins with a known function (Forgia et al., 2024).

The order *Tymovirales*, established in 2009, currently comprises five families: *Tymo-*, *Alphaflexi-*, *Betaflexi-*, *Gammaflexi-* and *Deltaflexiviridae*. While most members of the order infect plants, some infect fungi or insects (Adams et al., 2012a; Wang et al., 2012; Hamid et al., 2018; Kreuze et al., 2020). Members of the order *Tymovirales* typically have a 5.9-9.0 kb (+)RNA genome encoding a replicase in the 5’-proximal ORF, and express downstream genes via subgenomic RNAs (sgRNAs) (Adams et al., 2012b). Their genomes are 5’-m^7^G capped and 3’-polyadenylated (Adams et al., 2012b), except for viruses of the genus *Tymovirus*, which have a tRNA-like 3’ structure (Giegé et al., 1993). Three families of *Tymovirales* (*Alphaflexi-*, *Betaflexi-* and *Gammaflexiviridae*) have filamentous virions with lengths of 470-1000 nm and diameters of 12-13 nm, but virion morphology of the family *Tymoviridae* is icosahedral with a diameter of about 30 nm (Adams et al., 2012b). The family *Deltaflexiviridae* is thought to be capsidless because no virus particles have yet been found (Li et al., 2016a; Hamid et al., 2018; Dolja et al., 2020).

The family *Deltaflexiviridae*, established in 2017, currently has four recognized species (Li et al., 2016a; Marzano and Domier, 2016; Chen et al., 2016; Xiao et al., 2023), and prospective members of the family have been found in several agriculturally important fungi, such as *Botrytis cinerea*, *Sclerotinia sclerotiorum*, *Fusarium graminearum*, *Alternaria* spp. and *Pleurotus ostreatus* (Chen et al., 2016; Li et al., 2016a; Li et al., 2016b; Hamid et al., 2018; Ruiz-Padilla et al., 2021; Wang et al., 2022; Xiao et al., 2023). This family was defined based on the phylogeny of replication polyproteins, forming a monophyletic clade that is distinct from the other families of the order with high bootstrap value support, but its virion type and other properties remain largely unexplored.

In this study, we report isolation and characterization of a novel (+)RNA mycovirus, Fusarium oxysporum icosahedral virus 1 (FoIV1), from *Fusarium oxysporum* f. sp. *melonis*. Through phylogenetic analysis of their replication proteins, FoIV1 and three other, not all, previously identified deltaflexiviruses formed a monophyletic clade distinct from the other members of the family, suggesting that the clade could represent a new family. We also found that FoIV1 generates an sgRNA to express downstream small ORFs, of which ORF4 encodes an 18 kDa protein that constitutes the icosahedral capsids, as demonstrated by immunogold transmission electron microscopy. These results, as well as those obtained by clustering analyses and protein structure predictions, suggest that most deltaflexiviruses might encode a protein with a single jelly-roll (SJR)-like fold homologous to the ORF4 protein of FoIV1. These results provide clues to the evolutionary trajectory of viruses within the order *Tymovirales*.

## Materials and methods

### Fungal isolates and culture conditions

*F. oxysporum* f. sp. *melonis* strain Fom 405 (race 1, 2y) was isolated from a melon plant showing wilt in Ibaraki prefecture of Japan in 2006. Fom 405 and its virus-free isolates were cultured on PDA (24 g/L Difico Potato dextrose powder and 15 g/L agar) at 25°C for 7 days, and the mycelial discs were stored at -80°C in 25% glycerol. For collecting mycelia to extract total RNA and dsRNA, and to purify virus particles, mycelial discs were inoculated in PDB (24 g/L Difico Potato dextrose powder) and cultured with reciprocal shaking (60 strokes per min) at 25°C for one week.

### Isolation of virus-free isolates

The Fom 405 strain was cultured on PDA for about 2 weeks, and the surface of the colony was gently scraped and confirmed for production of microconidia under a microscope. The collected microconidia were suspended in 200 µl of sterile water, then the 10-fold serial dilutions of the suspension of the microconidia were plated on PDA and cultured for 3 days. Single colonies were picked and subcultured on PDA for 3 days, then the isolates were incubated in 50 ml PDB for one week. The presence of the virus was confirmed by viral dsRNA purification and specific reverse transcription PCR (RT-PCR) as described later.

### Determination of viral RNA sequence and genome properties

The viral dsRNA was purified from strain Fom 405 following the cellulase spin column method of Okada et al., 2015. Random-primed cDNA and products of rapid amplification of cDNA ends (RACE) of the dsRNA were cloned, Sanger-sequenced, and then assembled, as described previously (Okada et al., 2011; Okada et al., 2018).

The full-length viral sequence was subjected to ORF finder (https://www.ncbi.nlm.nih.gov/orffinder/) to determine the genome organization. The conserved domains of the hypothetical ORFs were analyzed using CD-search (https://www.ncbi.nlm.nih.gov/Structure/cdd/wrpsb.cgi) (Wang et al., 2023). For the hypothetical ORFs without any conserved domains, amino acid (aa) sequence-based pLM-BLAST (https://toolkit.tuebingen.mpg.de/tools/plmblast) (Kaminski et al., 2023) or HHpred (PDB_CIF70_8_Mar_database) (https://toolkit.tuebingen.mpg.de/tools/hhpred) (Zimmermann et al., 2018) was used to predict protein function.

### Phylogenetic analysis

Replicase-associated polyprotein sequences of viruses in the order *Tymovirales* were retrieved from Genbank (https://www.ncbi.nlm.nih.gov/genbank/) (Supplementary Table S1), and aligned with that of FoIV1 using MUSCLE5 (Edgar, 2022). The non-structural polyprotein sequence of Rubella virus (RUBV, NCBI Reference Sequence: NP_062883) was used as the outgroup. The alignments were submitted to IQ-TREE v1.6.12-stable release (Nguyen et al., 2015) with the option *-bb 1000*, MrBayes 3.2.7 (Huelsenbeck and Ronquist., 2001) on NGPhylogeny.fr online server (Lemoine et al., 2019) with default setting, and MEGA11 (Tamura et al., 2021) to produce maximum-likelihood (ML), Bayesian inference (BI) and neighbor-joining (NJ) trees, respectively, with default settings. The constructed trees were edited and visulized by Figtree v1.4.4 and Inkscape v1.3.2. In addition, SDT v1.3 program (Muhire et al., 2014) was used to calculate the pairwise identity of the replicase-associated polyprotein sequences of FoIV1, FoIV1-related viruses, and selected deltaflexiviruses (Supplementary Table S1).

### Plasmid construction

To obtain plasmids for preparing probes of northern hybridization, total RNA was extracted from 0.2 g of dried mycelium of the original Fom 405 strain using Trizol reagent (Thermo Fisher Scientific), following the manufacturer’s instructions. ORF1-ORF5 of FoIV1 were amplified by RT-PCR with specific primers (Supplementary Table S2) using SuperScript IV Reverse Transcriptase (Thermo Fisher Scientific) and GoTaq Master Mix (Promega), following the manufacturers’ instructions. The amplified fragments were cloned into pGEM-T Easy vector (Promega) and confirmed by Sanger sequencing to yield pGEM-ORF1-ORF5.

To construct a plasmid for protein expression, the ORF4 region was amplified by RT-PCR with specific primers (Supplementary Table S2) using PrimeScript One Step RT-PCR Kit Ver.2 (Takara Bio Inc., Kusatsu, Japan), following the manufacturer’s instructions, then cloned into pMD20 vector (Takara Bio Inc.), and confirmed by Sanger sequencing. The confirmed ORF4 fragment in pMD20 was digested with NdeI and XhoI and subcloned into the pET22b (+) vector (Merck) to yield pET-FoIV1-rORF4.

### Northern blot analysis

Ten micrograms of total RNAs of Fom 405 strain were denatured at 98°C for 5 min, then chilled on ice for 2 min. The heat-denatured RNAs were separated on a 1% agarose gel (95% 1× MOPS buffer and 5% formaldehyde) and transferred to a positively-charged nylon membrane (Roche) by capillary flow. After the blotting, the membrane was exposed twice to 120,000 μJ/cm^2^ for 1 min each time in a UV crosslinker (UVC500, Hoefer Inc., Holliston, MA). The membrane was then probed with the digoxygenin (DIG)-labeled DNA or RNA probes, and the hybridization signals were detected using the DIG detection kit (Roche) according to the manufacturer’s instructions. DNA probes were synthesized by PCR with DIG labeling kit (Roche), using template plasmids pGEM-ORF1-ORF5 and specific primers (Supplementary Table S2). Hybridization temperature of the DNA probes was 50°C. RNA probes complementary to positive-strand viral RNA genome were synthesized using DIG RNA labeling SP6/T7 kit combined with run-off transcription with linearized (SP6 promoter; *Nco*I digested) pGEM-ORF1-ORF5 as a template according to the manufacturer’s instructions. Hybridization temperature of RNA probes was 65°C.

### 5′ RNA ligase-mediated rapid amplification of cDNA ends (5′ RLM-RACE)

To detect sgRNAs, the 5’ RLM-RACE was performed with GeneRacer Kit (Thermo Fisher Scientific), using 1 μg total RNA of the original Fom 405 strain. The reverse transcriptions were performed with random primers according to the manufacturer’s instructions. Next, the primers ORF2-R2 and ORF2-R3 (Table S1), both targeting ORF2 region, paired with GeneRacer 5’ primer of GeneRacer Kit, were used for PCR reaction with KOD FX Neo (TOYOBO, Osaka, Japan) following the manufactures’ instructions. The amplicons were cloned into Zero Blun TOP vector and sequenced. The resultant sequences were aligned with the full-length FoIV1 genome using MegAlign software (Lasergene7, DNA-STAR).

### Purification and analysis of virions

All purification procedures were conducted at 4 °C, following the protocol in the published article (Howitt et al., 1995) with modifications. Twenty grams of dry mycelium was ground to a fine powder with liquid nitrogen and then mixed well with 100 ml buffer (0.1 M sodium phosphate, 0.2 M KCl, pH 7.4), added with 50 ml of chloroform, and vortexed until a homogeneous solution was formed. The solution was centrifuged at 11,000 × g (TOMY Suprema 21, NA-8 rotor, Japan) for 20 min, and the upper aqueous phase was ultracentrifuged at 120,000 × g (Hitachi CP80WX, P80AT rotor, Japan) for 90 min. The resulting pellet was resuspended in 3 ml of 0.1 M sodium phosphate buffer and gently rotated overnight. The suspension was centrifuged at 2,000 × g for 10 min, and the resulting supernatants were then ultracentrifuged at 120,000 × g (Hitachi CP80WX, P80AT rotor, Japan) for 90 min. The resultant pellet was resuspended with 1 ml of 0.1 M sodium phosphate buffer in a 1.5 ml tube overnight, and the suspension was centrifuged at 1,700 × g for 5 min to remove debris. The resultant supernatant was layered on the top of cesium chloride (CsCl) gradient (10-50%, 2 ml for each fraction and left overnight at room temperature) and centrifuged at 210,000 ×g (Hitachi 55P, RPS-40T rotor, Japan) for 2 h at 16°C. Six fractions were obtained using a gradient fractionator (HITACHI DGF-U, Japan) and then stored at -20°C before use.

The virus suspensions were analyzed by 13% SDS-PAGE followed by Coomassie Brilliant Blue (CBB) staining (EzStainAQua, ATTO, Japan). The observed 18 kDa protein bands (6.79 µg) were excised and subjected to the in-gel digestion and LC-MS/MS (LTQ XL, Thermo Fisher Scientific) analysis. The resultant peptide sequences were compared against the database of *Fusarium* spp.-related protein sequences (taxid: 5506) and predicted FoIV1-encoded proteins (accession number: BDQ13824-BDQ13828) using Mascot server v.2.4.1 (MatrixScience, Boston, MA).

To confirm the presence of viral RNAs in the virus suspensions, 50 µl suspension of each fraction was mixed with an equal volume of 2× STE buffer (20 mM Tris-HCl pH 8.0, 2 mM EDTA, 200 mM NaCl) containing 1% SDS, and 200 µl of phenol: chloroform: isoamyl alcohol (25:24:1). The mixture was vortexed for 10 min at room temperature, centrifuged at 15,000 × g for 5 min, and the aqueous phase was used as a template for RT-PCR with specific primer pair against FoIV1 ORF4 (Supplementary Table S2) using the SuperScript III One-Step RT-PCR System (Thermo Fisher Scientific).

### Antiserum production and immunoblotting

The pET-FoIV1-rORF4 plasmid was transformed into the BL21 Star (DE3) *E. coli* strain (Thermo Fisher Scientific) for protein expression. Transformed colonies were inoculated directly into 2× YT medium (1.6% polypeptone, 1% yeast extract, 0.5% NaCl, pH 7) supplemented with a defined volume of Overnight Express Autoinduction System 1 solution (Merck KGaA, Darmstadt, Germany). Cultures were incubated in baffled flasks at 37°C and 200 rpm overnight. The expression of FoIV1-rORF4 was induced in response to the growth of *E. coli*.

The bacterial cells were harvested by centrifugation at 3,000 × g for 10 min, and the wet weight of the cells was measured. For each gram of cells, 5 ml of BugBuster Protein Extraction Master Mix (Merck KGaA) was added, and the mixture was gently rotated at room temperature for 20 min to extract proteins. The pellet containing the FoIV1-rORF4 inclusion bodies was obtained by centrifugation at 12,000 × g for 20 min at 4°C and then suspended in 0.1× BugBuster Protein Extraction Reagent (Merck KGaA) using a homogenizer. The same process was repeated twice, and the purified inclusion bodies of FoIV1-rORF4 were pelleted by centrifugation at 12,000 × g for 20 min at 4°C, then solubilized by adding 8 M urea buffer (20 mM phosphate, 500 mM NaCl, 8 M urea, pH 7.4) containing 0.5% Triton X-100. After incubation at 37°C for 2 h, the supernatant obtained by centrifugation at 12,000 × g for 20 min at 4°C was applied to the column purification using immobilized metal affinity chromatography (IMAC) with a cobalt ion carrier (TALON metal affinity resin, Takara Bio Inc.). Antiserum was produced in a rabbit by immunization with the FoIV1-rORF4 (Eurofins Genomics).

For immunoblotting, the six fractions of the resulting CsCl gradient were analyzed by 13% SDS-PAGE, then transferred to PVDF membrane (ATTO, Japan). Proteins were detected using rabbit anti-FoIV1 rORF4-antibody (1:5000 dilution) as the primary antibody and HRP-conjugated goat anti-rabbit polyclonal antibody (Bio-Rad, #1706515, 1:10000 dilution) as the secondary antibody. Luminescent signals were detected using the EzWestLumi plus and EZ-Capture MG system (ATTO, Japan).

### Transmission electron microscopy (TEM)

The fraction of the purified virus suspension was negatively stained with 2% uranyl acetate (UA) and analyzed by TEM (JOEL 1400 Plus, JOEL, Japan) at 80 kV. Diameter of VLPs was measured by randomly selecting VLPs using ImageJ (Schneider et al., 2012) and calculating the average value.

Immunosorbent electron microscopy (ISEM) was performed following the previously published methods (Milne and Lesemann, 1984; Mueller et al., 2010) with slight modifications. Hydrophilic-treated carbon-coated copper grids (200 mesh, NISSHIN EM Co., Ltd, Tokyo, Japan) were placed on 20 µl droplets of rabbit anti-FoIV1 rORF4-antibody solution (1:1000 dilution in pH 7.4 phosphate-buffered saline, PBS buffer) for 15 min, then dried up with the filter paper. The grids were rinsed by placing on 20 µl droplets of PBS buffer for 5 min three times, then placed on 20 µl droplets of purified virus suspension for 15 min. The grids were rinsed by laying on 20 µl droplets of PBS buffer for 5 min three times and dried up on the filter paper. The samples were stained with 2% UA and analyzed by TEM.

Immunogold electron microscopy (IEM) was performed following the published article (Gulati et al., 2019). All the following procedures were performed in the humidity chamber. First, hydrophilic-treated carbon-coated copper grids were placed on 20 µl droplets of purified virus suspension for 2 min, then dried up with the filter paper and rinsed on 20 µl droplets of distilled water for 1 min. The grids were then placed on blocking buffer (pH7.4 PBS buffer, 0.3% BSA, and 0.1% Tween 20) for 20 min and dried up with the filter paper, then incubated on 20 µl droplets of rabbit anti-FoIV1 rORF4-antibody solution (1:2000 dilution in blocking buffer) for 90 min. The grids were rinsed by applying 20 µl droplets of wash buffer (pH 7.4 PBS buffer, 0.03% BSA, and 0.1% Tween 20) for 3 min, repeated five times, and then dried with the filter paper. The grids were then incubated on 20 µl droplets of gold-conjugated goat anti-rabbit IgG secondary antibody solution (5 nm gold, Cat#: EMGAR5, BBI solutions) for 60 min, then rinsed five times as above. Finally, the samples were stained with 2% UA, then dried up with the filter paper and analyzed by TEM.

### Protein sequence clustering analysis

To perform the coat protein (CP) analysis, we first fetched the domain annotations from the RVMT project (Neri et al., 2022), and extracted the amino acid regions for all CPs from the order *Tymovirales*. The potential CP ORFs of deltaflexiviruses and FoIV1-related viruses were screened using HHpred or pLM-BLAST. Overall, 379 sequences were extracted, which were then supplemented with the CP sequences of interest. Subsequently, we performed an “all vs all” pairwise alignment by using DIAMOND v2.0.15 (Buchfink et al., 2015) to search the protein set against itself (using diamond blastp with the flags --ultra-sensitive --id 15 --evalue 0.001 -k 1000000), followed by filtering the results by the alignments statistical estimates (alignment length ≥ 40aa, *E*-value ≤ 1□×□10^-3^) generating pairwise distance matrix which was then converted to a similarity network, visualized in Figure 4A, using Cytoscape v3.10.1 (Shannon et al., 2003), with the Prefuse Force-Directed layout, where edge weights are proportional to the -log(*E*-value). Independent of the node layout, connected components in the similarity network were identified and highlighted, and each connected component was then text labeled based on its protein annotations and phylogeny of the original sequence.

### Protein structure prediction and structure comparison

To predict the protein structure of CP of FoIV1 and its homologous proteins encoded by the selected deltaflexiviruses, we submitted protein sequences to the AlphaFold2/ColabFold v1.5.5 (Mirdita et al., 2022) with the default setting. Rank 1 prediction of each virus was then submitted to the DALI protein structure comparison server (Holm, 2022; Holm et al., 2023) to search the possible protein structures. The predicted structures were visualized using UCSF ChimeraX v1.7.1 (Pettersen et al., 2021).

The predicted CP structures of deltaflexiviruses and FoIV1 were subjected to the DALI server for all-against-all structure comparison. The dendrogram and heatmap were plotted following the published article (Butkovic et al., 2023a) using the R code from Dr. Anamarija Butkovic of Institut Pasteur, France, with the similarity matrix generated from the DALI server.

## Results

### New (+)RNA virus from a strain of *Fusarium oxysporum* f. sp. *melonis*

We extracted dsRNAs to identify viruses infecting a melon wilt fungus, *F. oxysporum* f. sp. *melonis* strain Fom 405 (Fig. 1A). A dsRNA segment of approximately 8.2 kbp was detected in 1% agarose gel electrophoresis (Fig. 1B). The 8.2 kb RNA sequence was determined by assembling the sequences of Sanger-sequenced cDNA and RACE clones. The length of the 5′ capped RNA segment is 8125 nt, excluding the 3′-terminal poly (A) tail (Fig. 1C). The sequence contains five ORFs. ORF1 encodes a putative viral replicase of 1962 aa (220 kDa) containing a methyltransferase, a helicase, and RNA-dependent RNA polymerase (RdRp) domains (Fig. 1C). ORF2, ORF3, ORF4, and ORF5 encode a 120 aa (13.2 kDa), 123 aa (14.3 kDa), 173 aa (18.2 kDa), and 142 aa (16.1 kDa) hypothetical proteins, respectively (Fig. 1C). The obtained viral sequence was deposited in Genbank of the National Center for Biotechnology Information (NCBI) with the accession number LC722819.

**Figure 1.**
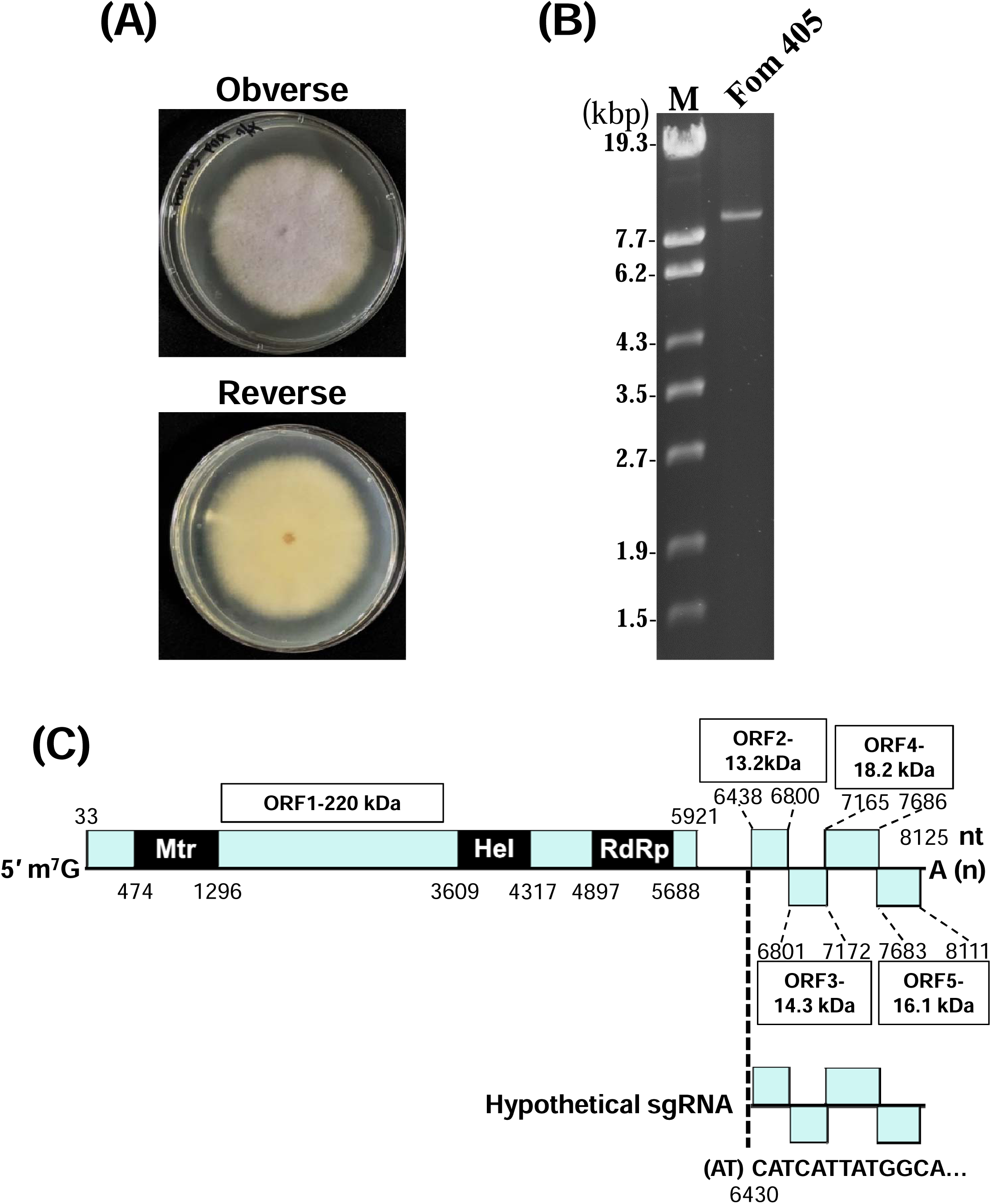
Novel RNA virus in *Fusarium oxysporum* f. sp. *melonis*. **(A)** Colony morphology of *F. oxysporum* f. sp. *melonis* strain Fom 405 cultured on PDA for 5 days at 25 °C. **(B)** dsRNA isolated from *F. oxysporum* f. sp. *melonis* strain Fom 405 was analyzed on a 1% agarose gel with EtBr (0.5 μg/ml) at 18 V for 20h. Lane M: 250 ng of λ-EcoT14I-digested DNA marker. **(C)** Schematic genome organization of FoIV1. Nucleotide sequence of the 5′ terminus of hypothetical sgRNA is shown according to the 5′ RLM-RACE result (Supplementary Fig. S3). ORFs are shown as boxes. ORF1: replicase-associated protein. Mtr, methyl transferase. Hel, helicase. RdRp, RNA-dependent RNA polymerase; ORF2, ORF3, ORF4, and ORF5 are hypothetical proteins.

A BLASTp search (*E*-value□≤□1□×□10^-5^) revealed that the ORF1-encoded protein is closely related to the replication proteins of viruses in the family *Deltaflexiviridae*. The closest related viruses are Calypogeia fissa associated deltaflexivirus (CfaDFV: Genbank accession: CAH2618741, 94% coverage, 64.4% identity), Erysiphe necator associated flexivirus 1 (Genbank accession: QKN22686, 95% coverage, 58.0% identity), Pestalotiopsis deltaflexivirus 1 (Genbank accession: QTH80200, 93% coverage, 57.5% identity) and Aspergillus flavus deltaflexivirus 1 (AfDFV1: Genbank accession: UAW09565, 97% coverage, 55.6% identity). In accordance with the current ICTV demarcation criteria for this lineage (Jan et al., 2017), a novel member is defined if it differs in host range, number of minor ORFs, or an amino acid identity (of the replication polyproteins) to known members below 70%. The virus isolated from *F. oxysporum* f. sp. *melonis* strain Fom 405 meets these criteria, and can thus be regarded as a novel species in the family. We named this novel virus ‘Fusarium oxysporum icosahedral virus 1 (FoIV1)’, due to the fact that, as described below, this virus has icosahedral virion.

Four FoIV1-free isolates (Fom 405-VF5, -VF15, -VF24 and -VF29) were obtained by conidial isolation that did not contain the 8.2 kbp dsRNA segment (Supplementary Fig. 1A), nor was the 522 bp-specific RT-PCR band detected (Supplementary Fig. 1B). After 7 days incubation on PDA, there was no significant difference in growth rates or phenotype between the four virus-free strains and virus-infected strains (Supplementary Fig. 1C).

### FoIV1 forms a phylogenetically distinct group within the order *Tymovirales*

Phylogenetic trees were constructed using ML (Fig. 2A), BI (Fig. 2B), and NJ (Fig. 2C) algorithms based on a multiple alignment of FoIV1 replicase-associated protein sequences with selected viruses in the order *Tymovirales* (Supplementary Table S1). The tree topologies obtained from all three methods are similar, with viruses in the family *Deltaflexiviridae* distinct from the other four families in the order. FoIV1 was placed in a distinct, monophyletic cluster (called ‘Group B’ hereafter) supported by high bootstrap values with three other viruses, CfaDFV, AfDFV1, and Lentinula edodes deltaflexivirus 1 (QOX06047, LeDFV1), in the ML tree (Fig. 2A, bootstrap value: 100%), BI tree (Fig, 2B, posterior probability: 1), and NJ tree (Fig, 2C, bootstrap value: 100%). FoIV1 shares 44-63% pairwise identity with the three members of the same group (Fig, 2D), and less than 30% with the other members of the family *Deltaflexiviridae* (called ‘Group A’ hereafter) (Fig. 2D). Multiple alignment of deltaflexivirus RdRp domain amino acid sequences showed that FoIV1 and the three related ‘Group B’ viruses all have canonical RdRp motifs A - C (Fig. 2E) (Urayama et al., 2024), but a notable number of amino acid residues within these motifs of the four viruses differ from those of other members of the family *Deltaflexiviridae*-GroupA (Fig. 2E). Specifically, motif C of Group B is NGDD[A/G] as opposed to SGDD[S/C/G/M] in the other viruses. Similarly, their motif A, DVTR WD[V/G]GCD, differs from DYTAWD[NSG]G[ICV]D). These results suggest that FoIV1, together with the three other viruses of the same cluster, belongs to a new viral family in the order *Tymovirales*.

**Figure 2.**
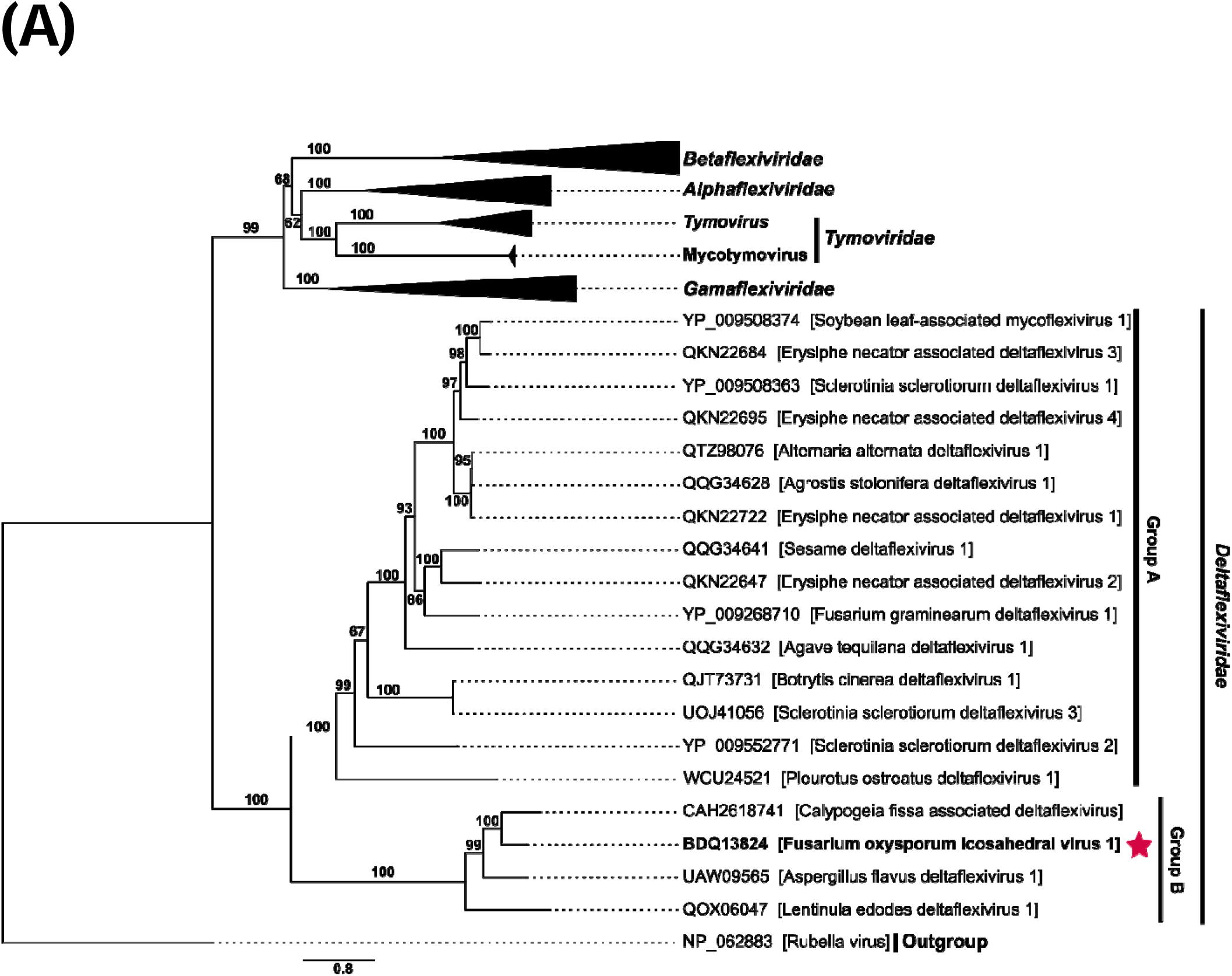
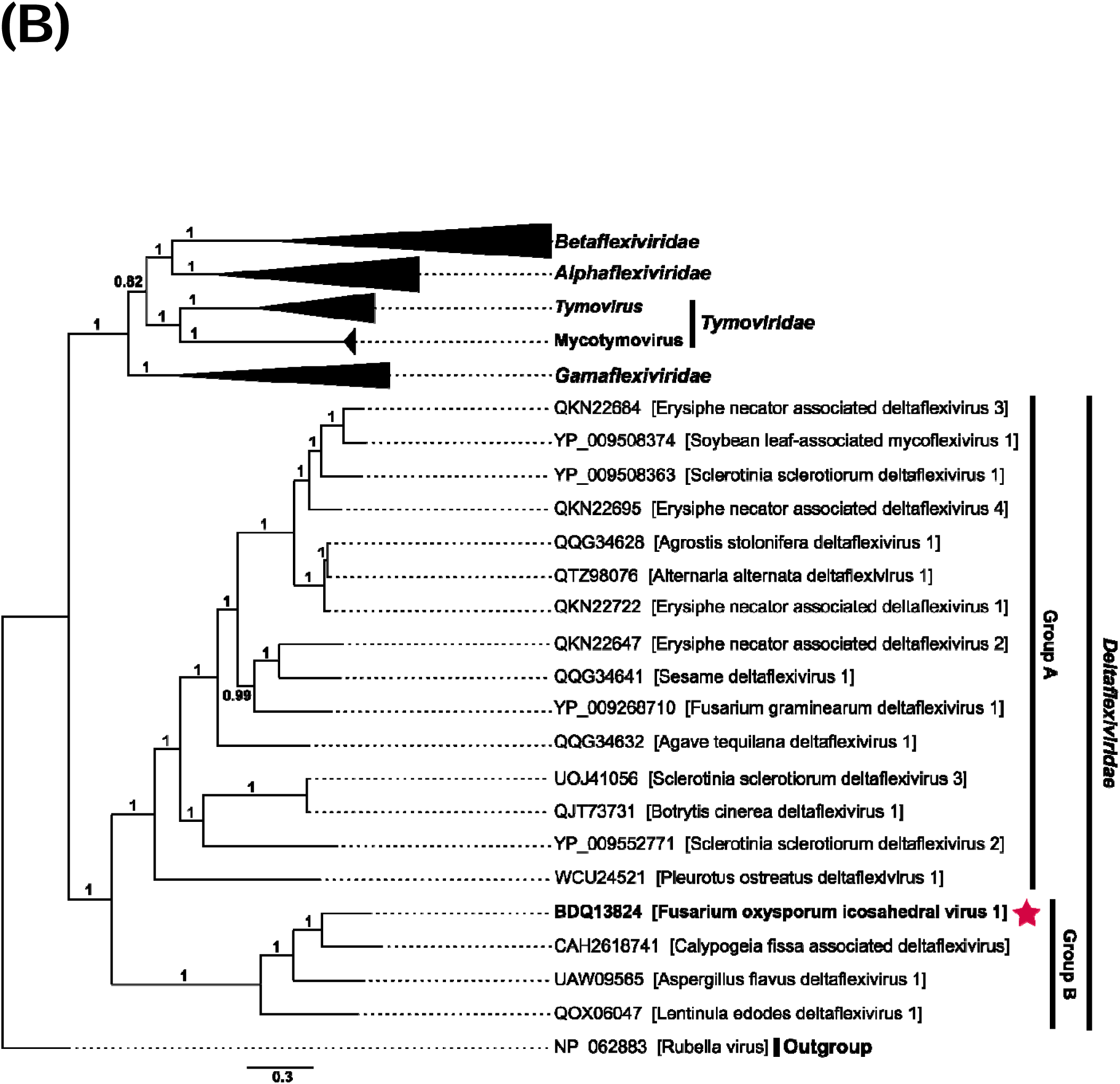
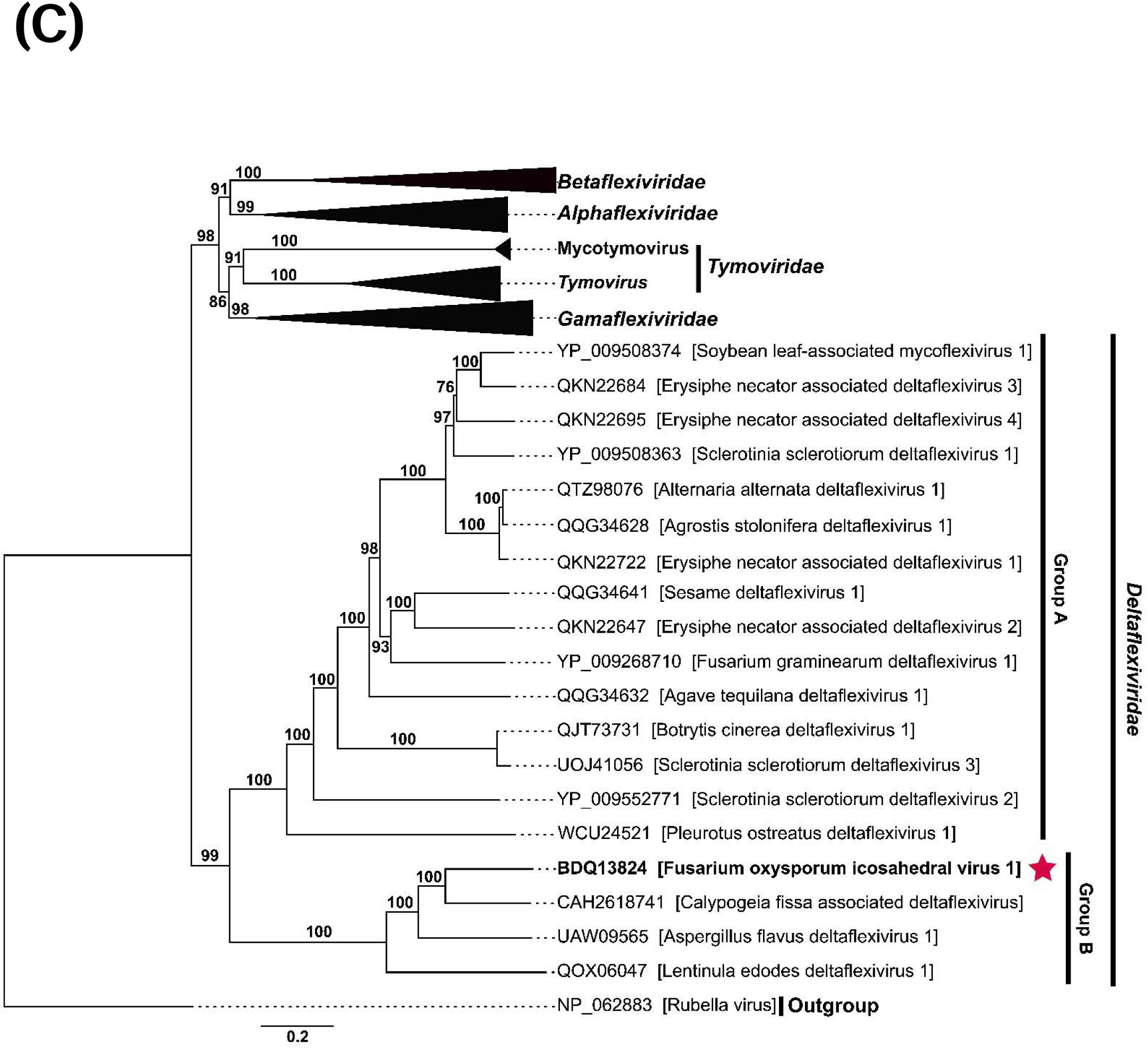
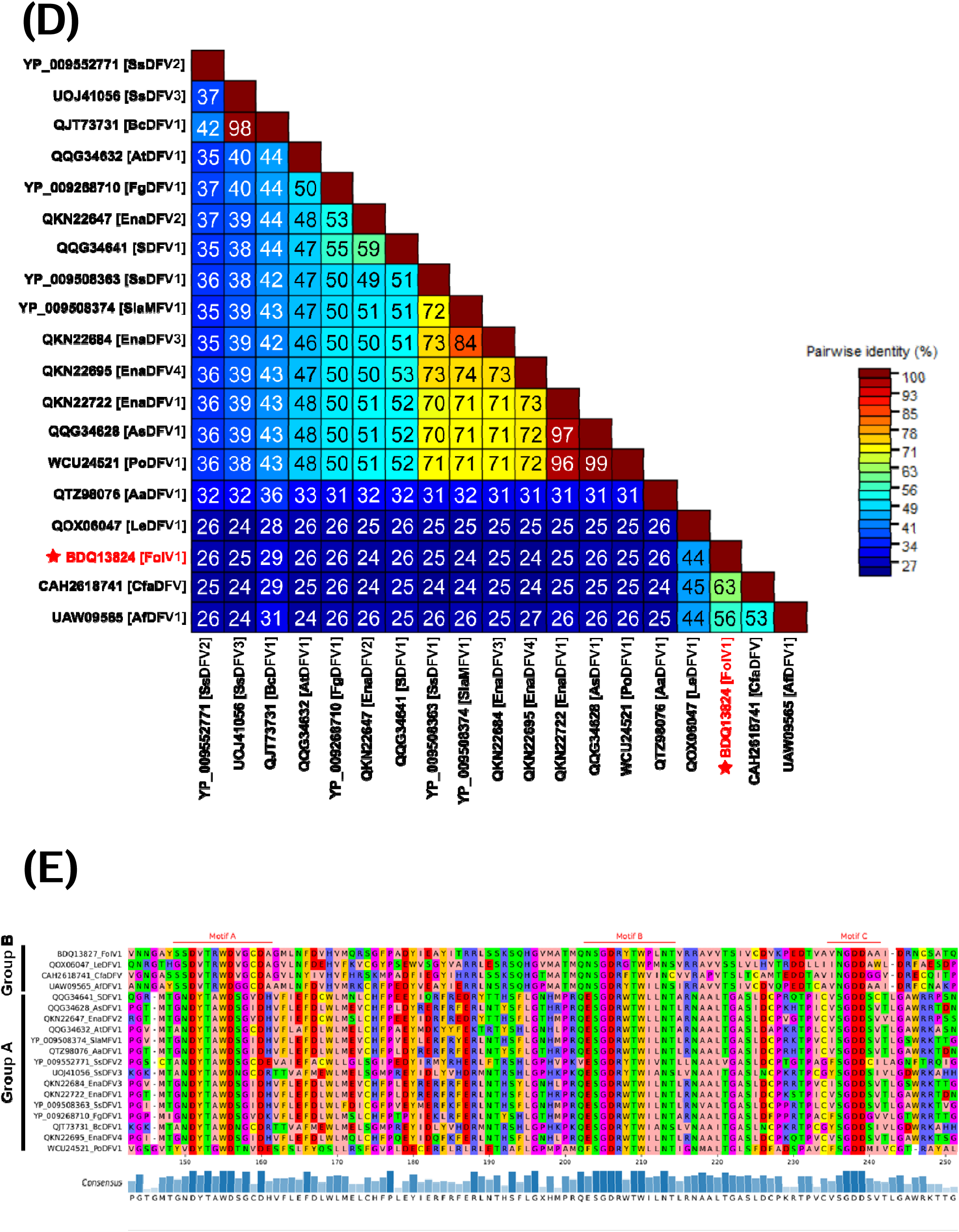
Phylogenetic analyses of FoIV1 and viruses in the order *Tymovirales* using the amino acid sequences of replicase-associated proteins. **(A)** A Maximum-Likelihood (ML) tree was constructed using IQ-TREE v1.6.12. The consensus tree was obtained from 1000 replicates with ultrafast bootstrap analysis (Best-fit model: LG+F+R6). Bootstrap values (%) are indicated next to the nodes. **(B)** A BI tree was constructed using MrBayesian 3.2.7 with 100,000 generations. Posterior probabilities are next to the nodes. **(C)** A NJ tree was constructed using MEGA11. The consensus tree was obtained from 1000 replicates. Bootstrap values (%) are marked next to the branches. **(D)** A pairwise identity matrix was generated using SDT v1.3. The numbers shown in the matrix are the pairwise identity (%). The RdRp sequence of FoIV1 identified in this study is highlighted in red. **(E)** Multiple alignment of RdRp aa sequences of FoIV1, FoIV1-related viruses, and deltaflexiviruses, visualized using pyMSAviz v0.4.2.

### Detection of the 5’ end of sgRNA for the expression of downstream ORFs

According to the current report of the International Committee on Taxonomy of Viruses (https://ictv.global/taxonomy/), most members of the order *Tymovirales* generate sgRNAs for translation of downstream proteins other than their replicase. However, protein expression strategies of Group A viruses, as well as Group B viruses including FoIV1 and its three related viruses, have not yet been investigated. We used Northern blot analysis (Supplementary Fig. 2) and 5’ RACE to test if FoIV1 possesses sgRNAs.

Northern blot analysis of fungal total RNA using DNA and RNA probes against ORF1-5 (Supplementary Fig. 2A) detected only a full-length genomic RNA of 8.2 kb (Supplementary Fig. 2B and 2C). We further performed 5′-RLM-RACE to identify the 5’ end of the sgRNA using a primer designed to target the ORF2 region of FoIV1. Sequencing of the RACE clones revealed that the 5′-end sequences of all 39 clones started at nt 6429 of the viral genome (Supplementary Fig. 3), although the first two nucleotides are adenine (A) and thymine (T), instead of T and guanine (G) that are found in the genomic sequence of FoIV1 (Figure S3). These results suggest that FoIV1 has an sgRNA for the expression of downstream small ORFs as a gene expression strategy (Fig. 1C), although it was below the detection limit of Northern blot analysis.

### FoIV1 ORF4-encoded protein is a component of the icosahedral virion capsid

No virions have been previously reported in deltaflexiviruses, and it remains unknown whether ORFs other than the replicase-encoding ORF1 encodes CP, because proteins encoded in this group of viruses do not share significant amino acid homologies with CP of other encapsidated viruses (Li et al., 2016a; Hamid et al., 2018). However, a search queried with the FoIV1 sequence of ORF4 using HHpred, which can predict protein functions by identifying homologous sequences with known protein structures, yielded five significant hits (96.1-97.3% probability) to the CPs of tymoviruses, including its type member turnip yellow mosaic virus (TYMV) (Table 1). The ORF2-, ORF3-, and ORF5-encoded proteins did not yield significant hits to any viral proteins (data not shown). This result suggests that FoIV1 has an icosahedral virion, similar to the tymoviruses.

**Table 1.** Result of HHpred using amino acid sequence of FoIV1 ORF4 as a query.

| No. | Hit | PDB ID | Probability (%) | Target length (amino acid) |
| --- | --- | --- | --- | --- |
| 1 | Citrus sudden death-associated virus; coat protein | 7SQZ | 97.29 | 136 |
| 2 | Desmodium yellow mottle virus; coat protein | 1DDL | 96.64 | 140 |
| 3 | Turnip yellow mosaic virus; coat protein | 2FZ2 | 96.55 | 140 |
| 4 | Citrus sudden death-associated virus; coat protein | 7SQY | 96.52 | 138 |
| 5 | Physalis mottle virus; coat protein | 1E57 | 96.13 | 146 |

To test this assumption, we tried to purify virus particles from fungal strain Fom 405 using CsCl gradient ultracentrifugation, using a modified method that used more starting material than previous studies (Howitt et al., 1995; Li et al., 2016a). There was a single approximately 18 kDa band of viral protein by SDS-PAGE in a suspension before CsCl gradient ultracentrifugation, and in fraction 2 obtained after centrifugation (Fig. 3A). Because the size of this band is similar to the estimated molecular mass of the protein encoded in ORF4 of FoIV1 (18.2 kDa), we produced an antibody against recombinant ORF4 protein (anti-FoIV1-rORF4) and performed immunoblotting with this antiserum. The anti-FoIV1-rORF4 reacted with this 18 kDa band, suggesting that the 18 kDa protein is the product encoded by ORF4 (Fig. 3B). In-gel digestion and LC-MS/MS analysis of the 18 kDa band yielded peptides derived from the estimated aa sequence of ORF4 of FoIV1 (Supplementary Fig. S4), confirming that the 18 kDa protein is the product of ORF4. Furthermore, FoIV1-specific RT-PCR detected the viral genome sequence of the ORF4 region in unfractionated suspension, and in fractions 1-3 from the CsCl density gradient (Fig. 3C), indicating the presence of viral nucleic acids associated with this 18 kDa protein. We performed TEM on the fraction 2, which contained both viral proteins and nucleic acids. First, by conventional TEM, we observed virus-like particles (VLPs, buoyant density: 1.26 g/cm^3^) with a diameter of approximately 32.5±4.2 nm (Fig. 3D). For further confirmation, we used anti-FoIV1-rORF4 antibody for ISEM and IEM. ISEM showed that the VLPs could be captured by the specific antibody coated on the copper grids (Fig. 3E). In the IEM analysis using grids absorbed with fraction 2, the gold particles clearly labeled the VLPs, confirming that the VLP was composed of ORF4 proteins (Fig. 3F). These results demonstrate that FoIV1 has icosahedral virions and that its structural protein, CP, is encoded by ORF4.

**Figure 3.**
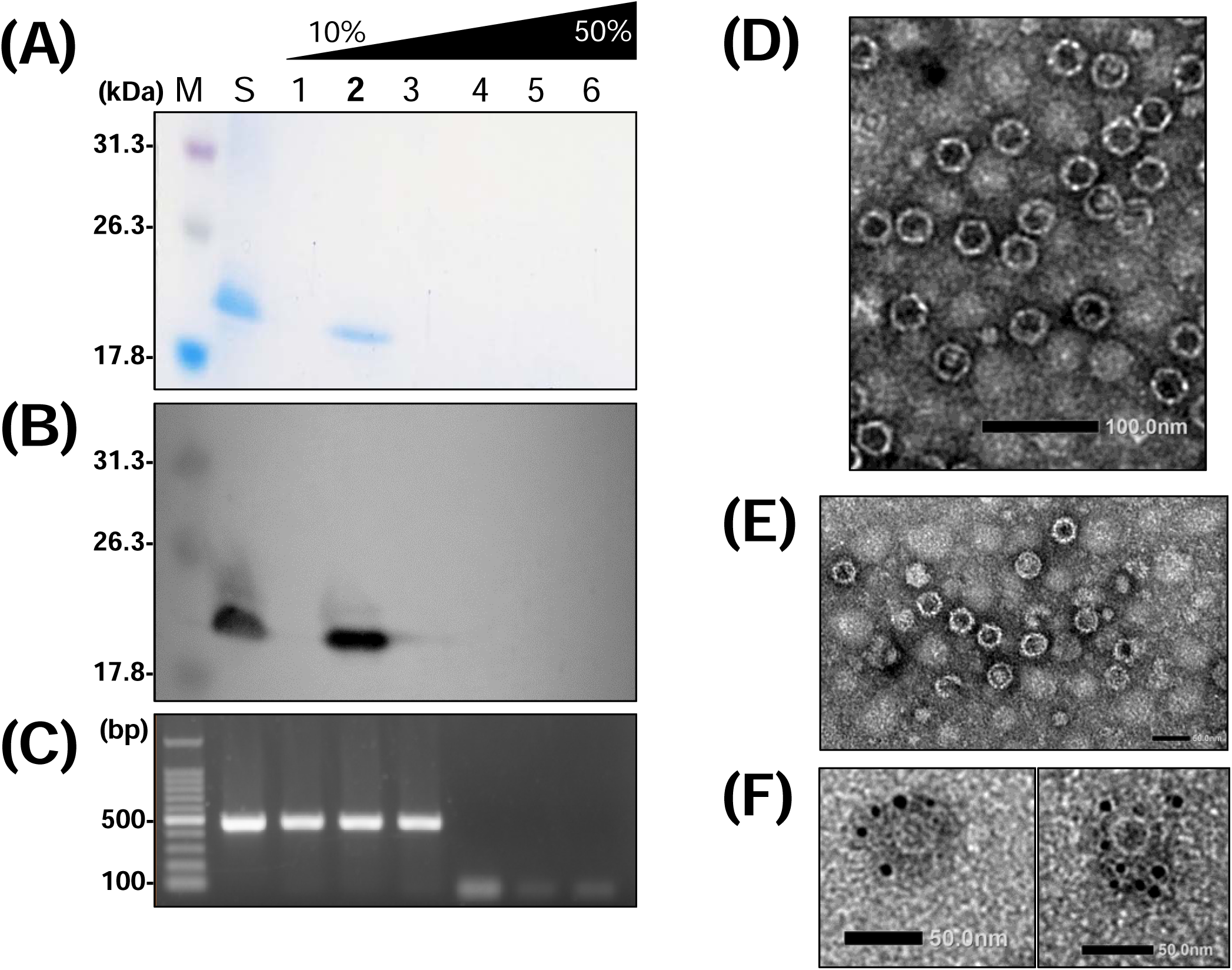
FoIV1 virus purification. Purified virus suspension was resolved with a 10-50% CsCl gradient, then fractioned into six fractions. Lane S, virus suspension before resolved in a CsCl gradient. **(A)** SDS-PAGE electrophoresis of the CsCl gradient fractions. The fractions were resolved in a 13% polyacrylamide gel at 120V for 2.5 h, then stained with CBB. **(B)** Western blot analysis of CsCl gradient fractions. Protein bands were detected with antiserum raised against FoIV1 ORF4-encoded protein. **(C)** Agarose gel electrophoresis of RT-PCR products using the primer pair ORF4-F and ORF4-R (Supplementary Table S2) against full-length FoIV1 ORF4. Fraction 2 showed an 18.2 kDa viral protein and viral nucleic acids, then was subjected to **(D)** conventional TEM, Scale bar: 100 nm. **(E)** ISEM, Scale bar: 60 nm. **(F)** IEM. Size of the gold particles: 5 nm. Scale bar: 50 nm. Virus particles are around 32 nm in diameter (buoyant density: 1.26 g/cm^3^).

### Deltaflexiviruses and FoIV1-related viruses encode potential SJR-like CP

Virion morphology in the order *Tymovirales* differs between families, which are either filamentous or icosahedral. In this study, we observed that the virions of FoIV1, which is closely related to viruses in the family *Deltaflexiviridae* based on replicase phylogeny, were icosahedral. To further examine the virion structures of deltaflexiviruses and viruses of the Group B that includes FoIV1, we screened potential CP-encoding ORFs using HHpred or pLM-BLAST. Searches against 15 viruses in the family *Deltaflexiviridae*-Group A and the three viruses of the family *Deltaflexiviridae*-Group B queried against FoIV1 CP yielded 14 significant hits (HHpred: Probability≤85%; pLM-BLAST: *E-value*□≤□1□×□10^-3^). However, based on the threshold value, we did not find any ORFs predicted to encode CP in any of the three deltaflexiviruses (SsDFV2, SsDFV3, and BcDFV1) or one virus (LeDFV1) of the family *Deltaflexiviridae*-Group B. Using these 14 potential CPs (Supplementary Table S3) and the ORF4 product of FoIV1, we performed a clustering analysis of the potential CP amino acid sequences with those of 397 mostly environmental viruses in the order *Tymovirales* (Fig. 4A), and predicted their protein structure by AlphaFold2/ColabFold (Fig. 4B). The predicted protein structures were subjected to the all-against-all structure comparison including CPs of TYMV (SJR, PDB ID: 2FZ2) and potato virus X (PVX, Phlebo NC-like, PDB ID: 6R7G), which has filamentous virions.

**Figure 4.**
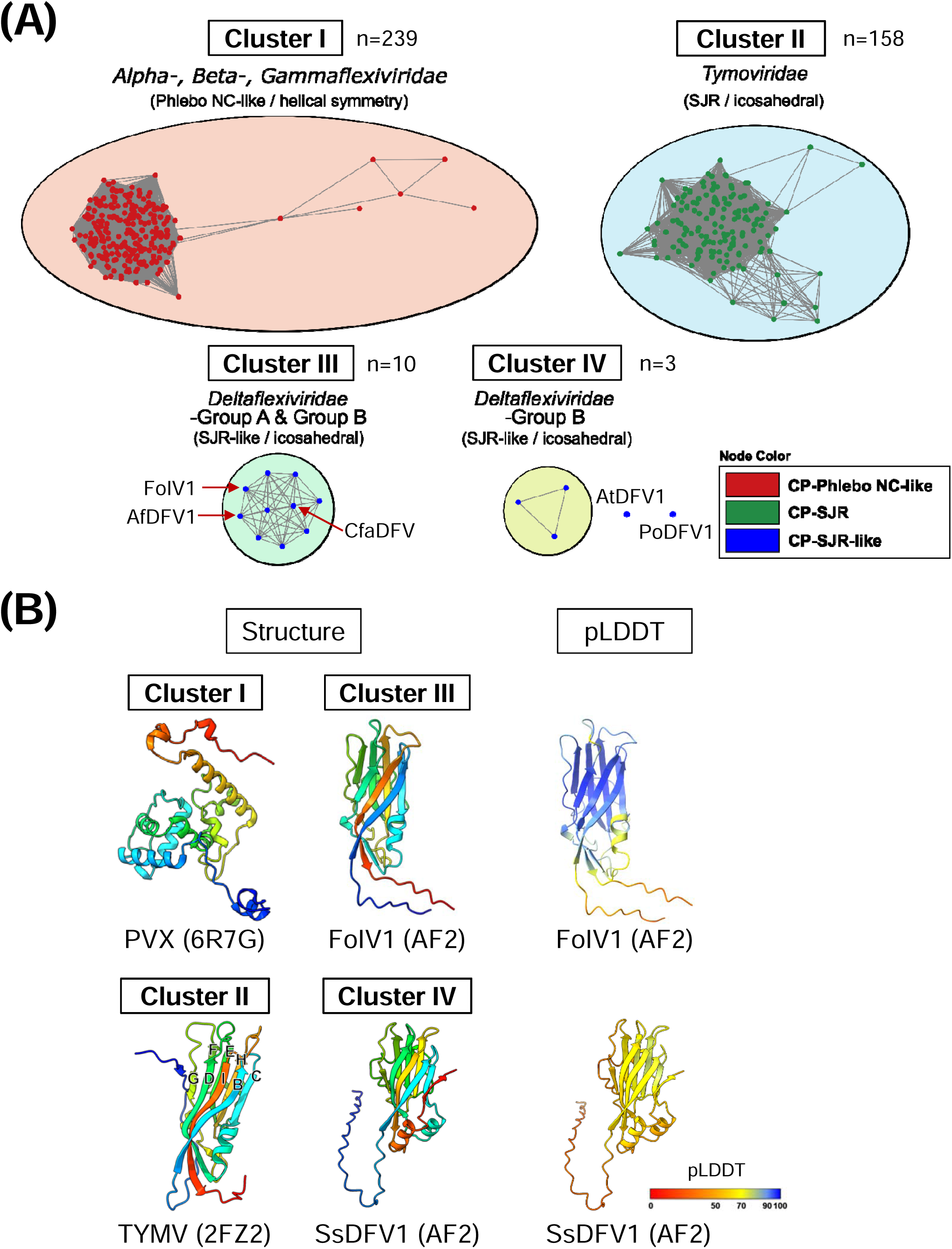
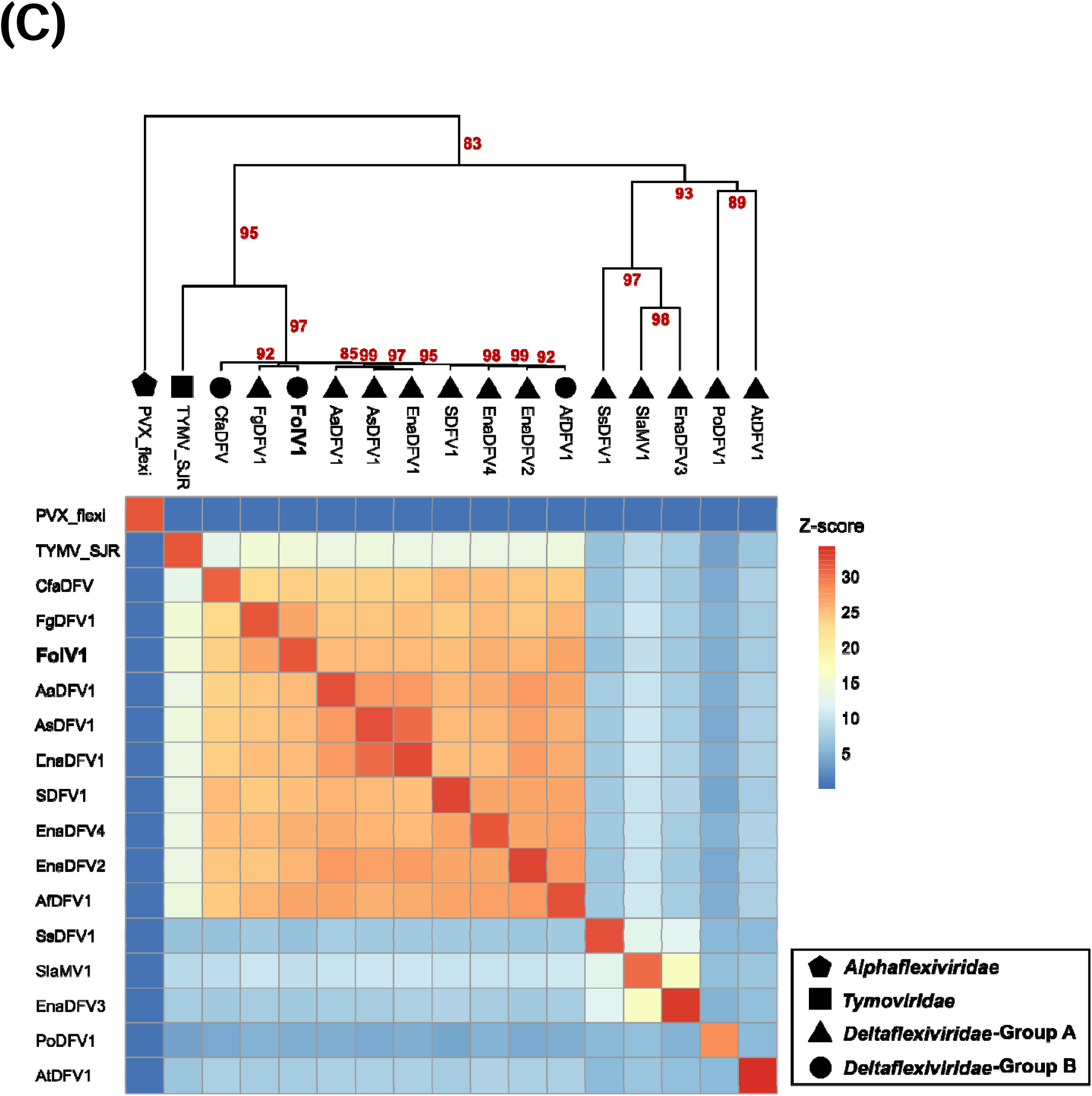
Sequence- and structure-based similarity analyses of the CPs of FoIV1, FoIV1-related viruses, and other members of the *Tymovirales*. **(A)** Sequence-based clustering of CPs of the *Tymovirales*. CP sequences were clustered using all vs all pairwise alignment of DIAMOND v2.0.15 (*E*-value ≤ 0.001), then visualized by Cytoscape v3.10.1. Four distinct clusters and two independent nodes are present. Numbers of nodes (n) are marked under the clusters. Nodes are colored by their protein folding types. The red arrows indicate the CPs of FoIV1-related viruses. **(B)** Protein structure modeling of CPs. Structures are color-coded with a rainbow color scheme from N-terminal (blue) to C-terminal (red). The pLDDT scores of Alphafold2 (AF2) predicted proteins are also shown in the figure. BIDG-CHEF β-strands of SJR folding structures are annotated on the model of TYMV CP. **(C)** Dendrogram and heatmap of all-against-all structure comparison of SJR CP of TYMV, Phlebo NC-like CP of PVX, and CPs of FoIV1-related viruses and deltaflexiviruses. The “Average” method was used for clustering (clustering methods: ward.D, ward.D2, single, complete, average, mcquitty, median, and centroid were tested to find the best clustering). Bootstrap values are indicated as red numbers at the nodes.

Sequence clustering separated *Tymovirales* CP sequences into four clusters and two independent nodes. *Alpha-*, *Beta-*, and *Gammaflexiviridae* with filamentous particles formed a single cluster (Fig. 5A, upper left, red nodes). Another distinct cluster solely contained CP sequences from members of the family *Tymoviridae*, which are assembled into icosahedral virions by SJR proteins (Fig. 4A, upper right, green nodes). Additionally, two other separate clusters and two nodes contained the potential CP sequences of deltaflexiviruses and *Deltaflexiviridae*-Group B, including the CP sequence of FoIV1 (Fig. 4A, blue nodes). However, the deltaflexivirus CP sequence clustering pattern is incongruent with its replicase phylogeny; the 12 deltaflexiviruses in the Group A, which is distinct from a clade containing the viruses of the family *Deltaflexiviridae*-Group B, in the replicase phylogeny separated into two clusters and two independent nodes (Fig. 2). This separation of SJR-type CP in the order *Tymovirales* into clusters II through IV was supported by their predicted CP structures (Fig. 4B and Supplementary Fig. S5). Whereas the canonical SJR-CP, including that of TYMV, has eight β-strands forming two antiparallel sheets (Cheng and Brooks., 2013), the predicted structures of putative CPs in the cluster III, which includes three *Deltaflexiviridae*-Group B viruses (Supplementary Fig. S5, represented as “*Deltaflexiviridae*-Group B”) and seven deltaflexiviruses (Supplementary Fig. S5, outside the green dashed box in the blue box of the family *Deltaflexiviridae*-Group A) have seven β-strands and a coil, with the antiparallel features of their N- and C-termini are similar to the canonical SJR-CP of cluster II. The other five deltaflexiviruses in the cluster IV (Supplementary Fig. S5, represented in a green dashed box) have eight β-strands, but their N-terminal coils are extremely long. Amino acid-based clustering and structure predictions suggest that SJR-like CPs of viruses in the conventional family *Deltaflexiviruses* can thus also be separated into two groups.

To further examine the structural relationships between the SJR-CPs of viruses in the *Deltaflexiviridae* and viruses in the *Tymoviridae*, we developed a matrix and cluster dendrogram based on pairwise CP structure comparisons obtained from DALI *Z*-scores. The CPs of FoIV1 and two Group B viruses, as well as seven Group A deltaflexiviruses, have *Z*-scores of 3.6-14.9 (*Z*-score 8-20 represents high probability of being homologous and 2-8 indicates a ‘gray area’; Benton and Himmel, 2023) with the SJR-CP of TYMV, and reside in the same cluster as TYMV CP, but not with the Phlebo NC-like CP of PVX, which belongs to the family *Alphaflexiviridae* (Fig. 4C). A distinct subcluster containing putative CPs of five deltaflexiviruses, SsDFV1, SlaMV1, EnaDFV3, PoDFV1, and AtDFV1 was found within the large SJR-CP cluster, and each structure has *Z*-scores of 3.6-8.9 with the SJR-CP of TYMV, which are lower than between TYMV CP and the other ten virus CPs. The results of the structure comparison are consistent with the sequence clustering analysis, in that potential CP sequences of these five viruses are excluded from both the *Tymoviridae* cluster (upper right in Fig. 4A) and the FoIV1-containing cluster (lower left in Fig. 4A). These results, along with the structure predictions, suggest that *Deltaflexi*-related viral particles are not filamentous particles like the *Alpha-*, *Beta-*, and *Gammaflexiviridae*, but are likely icosahedral like FoIV1.

## Discussion

The family *Deltaflexiviridae* was established based on replicase phylogeny and named to keep continuity with existing family names, *Alpha-*, *Beta*-, and *Gammaflexiviridae*, without considering properties such as gene expression strategies and the functions of encoded proteins. In this study, we report a novel (+)RNA mycovirus, FoIV1, and propose that this virus constitutes a new family within the order *Tymovirales* along with 3 previously reported deltaflexiviruses, as these viruses form a distinct sister clade branching outside other deltaflexiviruses with high bootstrap value support (Fig. 2A-C). Remarkably, ORF4 of FoIV1 encodes an SJR-like CP that assembles into icosahedral virions, contrary to the assumption that viruses in the “*-flexividae*” families are exclusively filamentous or non-encapsidated. By identifying a likely CP-encoding ORF and through structural comparisons, we demonstrate conclusively that many other deltaflexiviruses also have SJR-like CPs, implying that the family name—*Delta“flexi”viridae*, is not appropriate.

The most important result of this study is that we succeeded in purifying virions of FoIV1 and demonstrated that they have icosahedral capsids (Fig. 3). To our knowledge, this is the first report of the direct observation of deltaflexivirus-related virions, possibly due to a slightly modified virus purification protocol (Howitt et al., 1995); we doubled the amount of mycelium from 10 g to 20 g to contain more virions in the starting material. Additionally, post-ultracentrifugation pellets were suspended overnight so that the virions could fully dissociate. The most recent study that attempted to purify virions of deltaflexiviruses, SsDFV1, failed to observe virions, but the detailed procedures and conditions of purification were not explained (Li et al., 2016a). Since then, there have been no more published attempts to purify deltaflexiviruses virus particles, with the result that deltaflexiviruses were thought to be capsidless (Li et al., 2016a; Hamid et al., 2018). This conclusion was reasonable because no viral encoded proteins of deltaflexiviruses share significant amino acid homology with any known CPs of other viruses. However, protein structure predictions (Fig. 4B) and protein structure similarity analysis (Fig. 4C) demonstrated that the predicted structure of a protein encoded by ORF4 of FoIV1 is similar to the SJR-CP of tymoviruses, and the SJR-like CPs of FoIV1 and nine other viruses (seven Group A deltaflexiviruses and two FoIV1-related Group B viruses) clustered with TYMV CP. Although the predicted structure of FoIV1 CP was slightly different from the canonical SJR-CPs in that the β-strand “G” of SJR is replaced with a coil, its overall structure is similar to that of the canonical SJR fold. The Alphafold algorithm can help to predict protein structures, but Cryogenic electron microscopy (Cryo-EM) is needed to determine whether FoIV1 CP has seven instead of eight β-strands, and how they assemble into particles. Detailed structural information of FoIV1 CP will help to explore novel motifs involved in virus-host interactions and viral RNA packaging, as well as differences between the CPs of plant tymoviruses and FoIV1.

Purified FoIV1 virions were analyzed by immunoblotting (Figures 3A and 3B) and observed by immuno-EM (Figures 3E and F), which confirmed that FoIV1 ORF4 encodes CP, and also indicated that FoIV1 uses a certain strategy to express proteins encoded by small ORFs located downstream of the ORF encoding the replicase. (+)RNA viruses usually have at least one gene expression strategy, such as sgRNAs, internal translation initiation, leaky scanning, frameshift, or readthrough (Schirawski et al. 2000). Most members of the order *Tymovirales* employ sgRNAs to express downstream genes and have promoter sequences upstream of the sgRNAs (Komatsu et al., 2012; Kreuze et al., 2020; Schirawski et al. et al., 2000). Using 5’ RLM-RACE (Figure 1C, Supplementary Fig. S3), we found that FoIV1 has only one sgRNA, whose transcription starts upstream of ORF2 but not ORF4, and no subgenomic promoter sequences were predicted upstream of the transcription start site of ORF2. Because the sgRNA has a short 5′ untranslated region (9 nt), it is possible that FoIV1 ORF4 was translated by leaky scanning of upstream initiation codons in the sgRNA, as in the case of the potexviruses (Fujimoto et al., 2022).

Because we experimentally verified that the ORF4-encoded SJR-like CP of FoIV1 is a component of its icosahedral virions, at least nine viruses that cluster together in the pairwise CP structure comparisons are also likely to have icosahedral virions. However, we do not have enough evidence to speculate about the other five deltaflexiviruses whose CPs did not cluster with FoIV1, because the N-terminal coils of their predicted structures are much longer than the 10 SJR-like CPs. Given that HHpred or pLM-blast predictions suggest that they may be viral CPs (Supplementary Table S3), and that structurally unrelated proteins can form icosahedral virus particles (Krupovic and Koonin, 2017), it is possible that these five viruses also have icosahedral virions. Future attempts to purify viral particles of these viruses are needed to confirm the presence of virions and their morphology.

The evolutionary trajectory of CP genes in deltaflexiviruses remains enigmatic. FoIV1-related Group B viruses and Group A deltaflexiviruses belong to the same *Tymovirales* order as other mycoviruses, including gammaflexiviruses (Howitt et al., 2001) and mycotymoviruses (Li et al., 2016b), and members of each of these groups share similar genomic features with plant flexiviruses and plant tymoviruses, such as the 5’ proximal large ORF1 encoding replicase, conservation of replicase domains, and the use of sgRNA for downstream gene expression. However, the number of ORFs located downstream of the ORF1 can vary among these viruses, and ORF4 does not consistently encode CP (Supplementary Table S3). Furthermore, no CP-encoding ORF was found in some viruses within these groups. In fact, our results showed that four viruses (SsDFV2, SsDFV3, BcDFV1, and LeDFV1) are not predicted to have CP-encoding ORFs, and that the presence or absence of CP ORF is inconsistent with phylogeny based on the replicase. The most plausible explanation for this inconsistency is the loss of the CP-encoding gene in some mycoviruses. The study by Andika and colleagues provides evidence that certain viruses can infect both plant and fungal hosts, suggesting the possibility of host shifts between these kingdoms (Andika et al. 2017). Based on phylogenetic analyses of the order *Tymovirales* (Fig. 2), it is convincing that the ancestral virus of the deltaflexiviruses used plants as its primordial host (Martelli et al., 2007). Considering these lines of evidence, we can hypothesize that some fungal viruses lost their CP genes, since the CP is not always essential for mycoviruses, as exemplified by the hypoviruses (Suzuki et al., 2018) and endornaviruses (Valverde et al., 2019). In concordance with the reductive evolution of the CP in Group A and B deltaflexiviruses, the FoIV1 CP is small (18 kDa) compared with 20 kDa TYMV CP. The SJR-CP plays a critical role in mediating virus-host interactions, as suggested by a recent study showing that the viral SJR-CP can evolve to accommodate specific interactions with hosts, potentially leading to changes in its size and structure (Butkovic et al., 2023b). Collectively, the evidence suggests that the deltaflexiviruses CP is undergoing a reductive evolutionary process, possibly driven by adaptation to fungal hosts (Supplementary Fig. S6). Future analyses are needed to assess the biological significance and functional implications of the small CP of FoIV1 in the context of its fungal host. Such studies could provide insights into the molecular mechanisms underlying host adaptation and the evolutionary trajectories of these viral proteins. The identification of ORF4 encoding the FoIV1 CP provides an important clue to this process.

Initially, we would not have been able to determine if FoIV1 encoded CP by conventional BLAST or CD-searches. Nowadays, prediction of protein structure and clustering of proteins based on their structures is emerging as a powerful method that can efficiently estimate protein functions and help explore evolutionary processes of their encoding genes (Barrio-Hernandez et al., 2023; Butkovic et al., 2023a). In this study, protein homology search tools (HHpred and pLM-BLAST) combined with clustering provided essential clues for the discovery of FoIV1 virions and CPs, and evidence for the possibility that other deltaflexiviruses are encapsidated. In conclusion, this study extends our understanding of diversity within the order *Tymovirales*. The establishment of a new viral family to accommodate FoIV1 and its related viruses will facilitate more accurate and reliable classification for further research into the evolution and biology of viruses within the order *Tymovirales*.

## Data availability

The complete genome of FoIV1 was deposited in GenBank of NCBI with the accession number LC722819.

## Supplementary data

Supplementary data are available at *Virus Evolution* online.

## Acknowledgments

We would like to thank Dr. Syun-ichi Urayama (University of Tsukuba) for critical reading of the manuscript; Dr. Hiroshi X. Chiura (Tokyo University of Agriculture and Technology), Dr. Nobumitsu Sasaki (Tokyo University of Agriculture and Technology), Dr. Yuh-Kun Chen (National Chung-Hsing University), and Dr. Che-Yen Joseph Wang (Penn State University) for the fruitful discussions on virion purification and analyses; Dr. Yoshiyuki Ito (Tokyo University of Agriculture and Technology) and Mrs. Miki Hisada for helping on LC-MS/MS analysis; Dr. Anamarija Butkovic (Institut Pasteur) for sharing the codes for generating dendrogram and heatmap.

## Author contributions

C.-F.W., R.O., K.K, and H.M. designed the research; C.-F.W., R.O., Y.-C.C., O.T., and K.K. performed experiments; R.O. and O.T. contributed new materials; C.-F.W. and U.N. performed bioinformatics; C.-F.W., R.O., U.N., Y.-C.C., K.K., H.M., and K.K. analyzed the data; and C.-F.W., R.O., U.N., H.M., and K.K. wrote the manuscript.

---

**Supplementary Figure S1.**
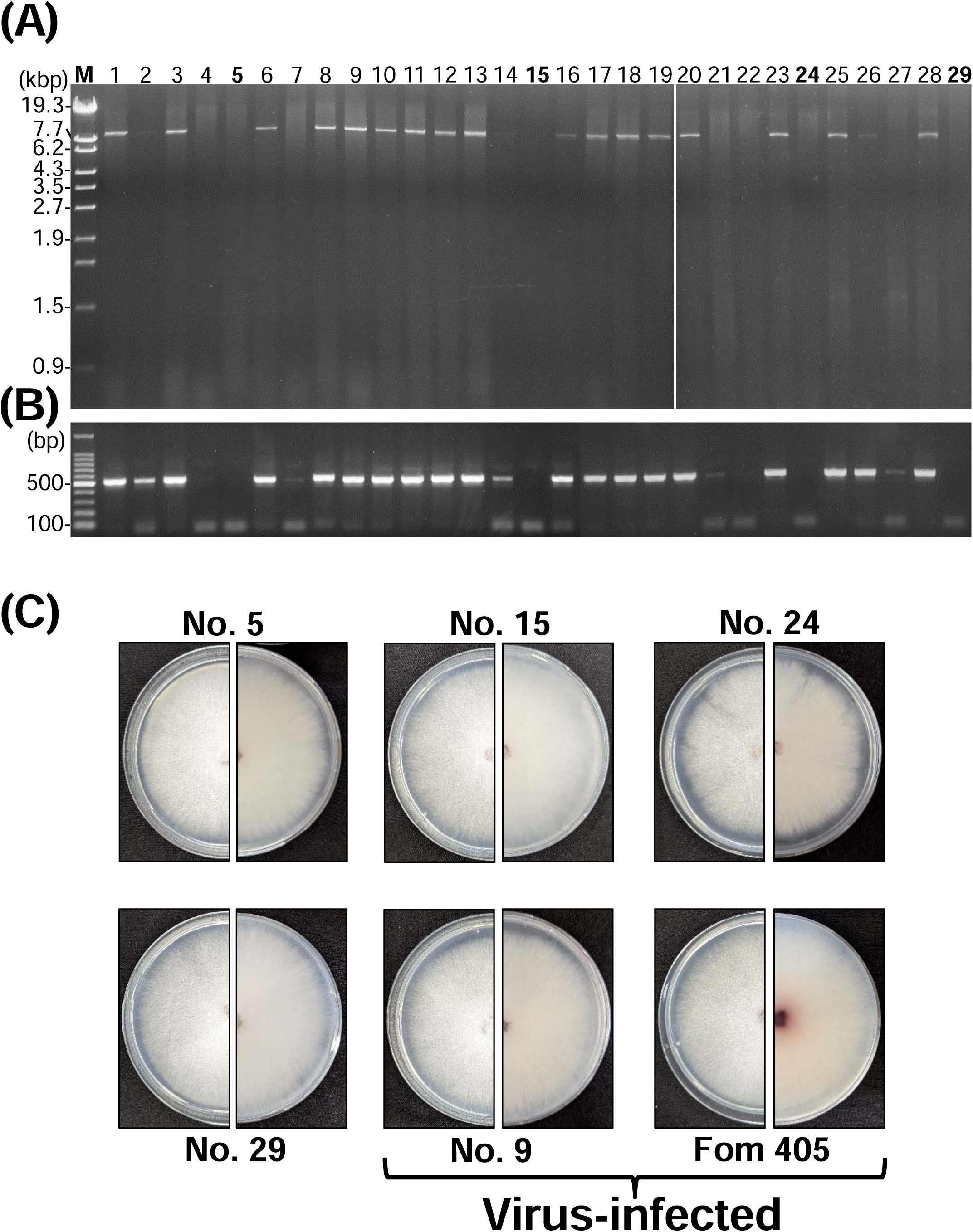
Obtainment of the FoIV1-free isolates by conidia isolation. **(A)** The dsRNAs were extracted from each conidial isolate and analyzed on a 1% agarose gel with EtBr (0.5 μg/ml) at 18 V for 20h. Lane M: 250 ng of λ-EcoT14I-digested DNA marker. The numbers in bold are FoIV1-free isolates. **(B)** No specific FoIV1 RT-PCR band (521 bp) was detected from isolates No.5, No.15, No.24, and No.29. **(C)** Colony morphologies of FoIV1-free and virus-infected isolates.

**Supplementary Figure S2.**
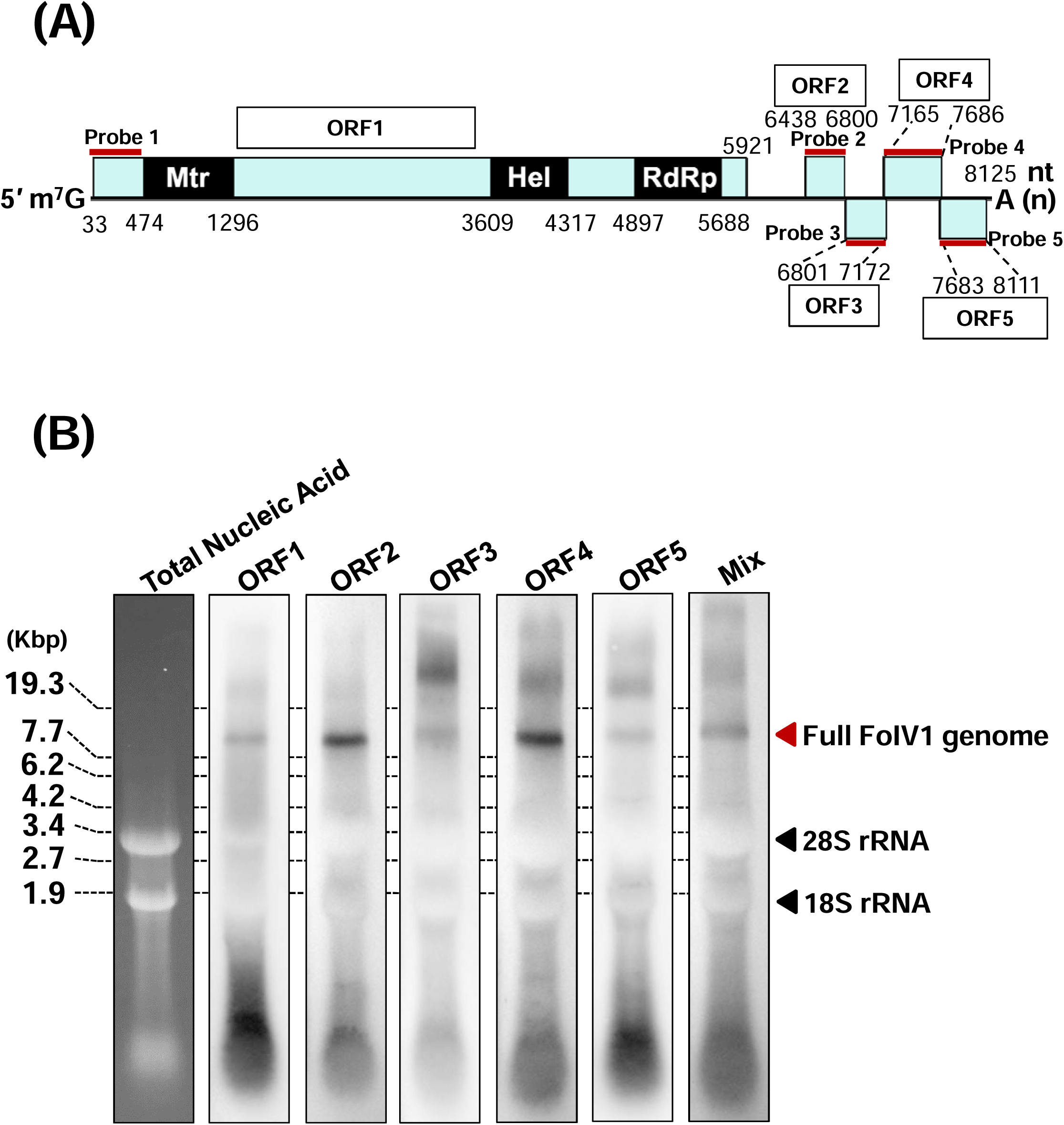
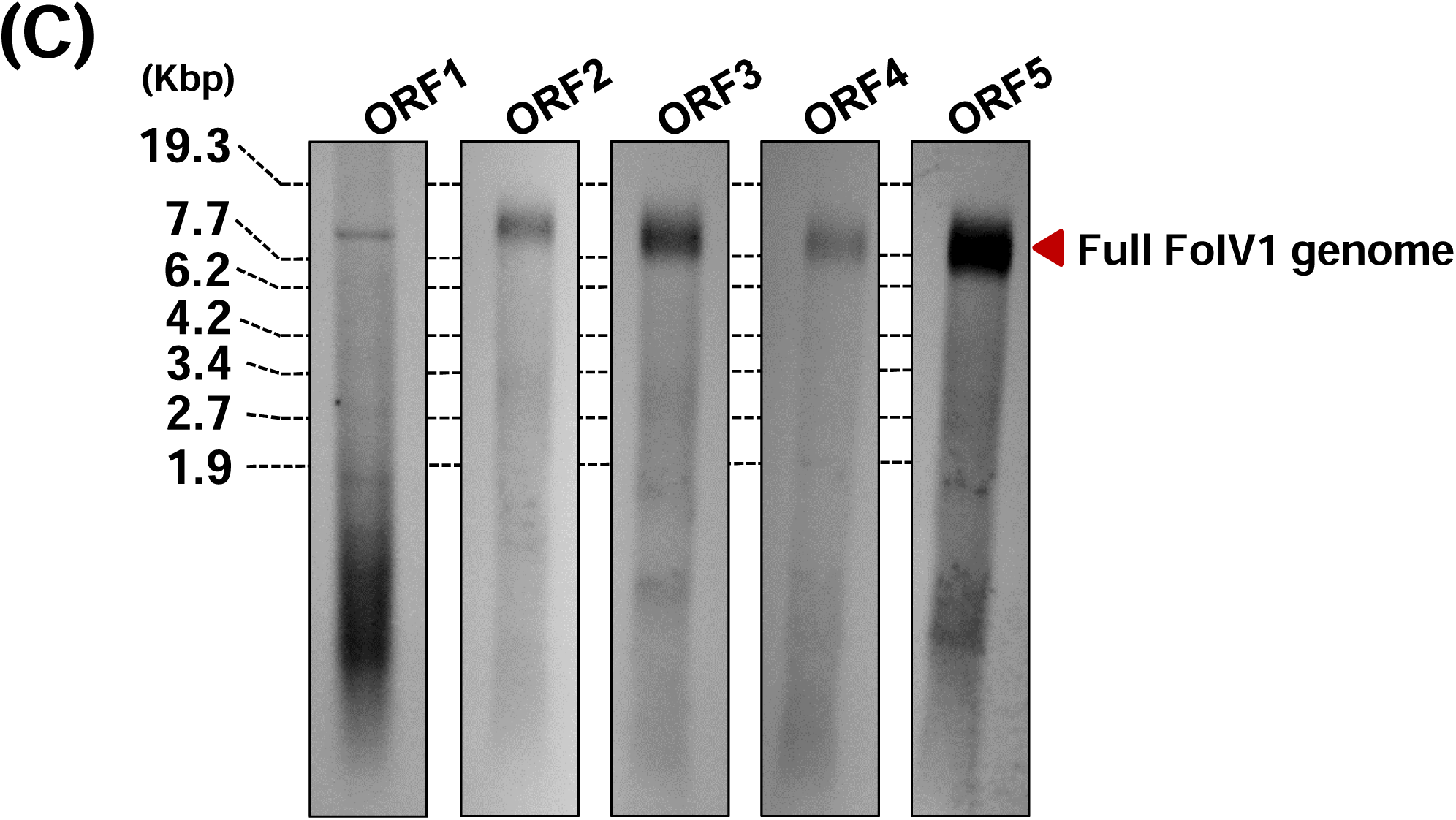
Detection of the FoIV1 sgRNA using northern hybridizations. **(A)** Probe positions of **(B)** DIG DNA probe hybridization and **(C)** DIG RNA probe hybridization. Probes are shown as red bars. ORFs are shown as boxes. Position of the full-length FoIV1 genome (8.2 kb) is indicated at the right.

**Supplementary Figure S3.**
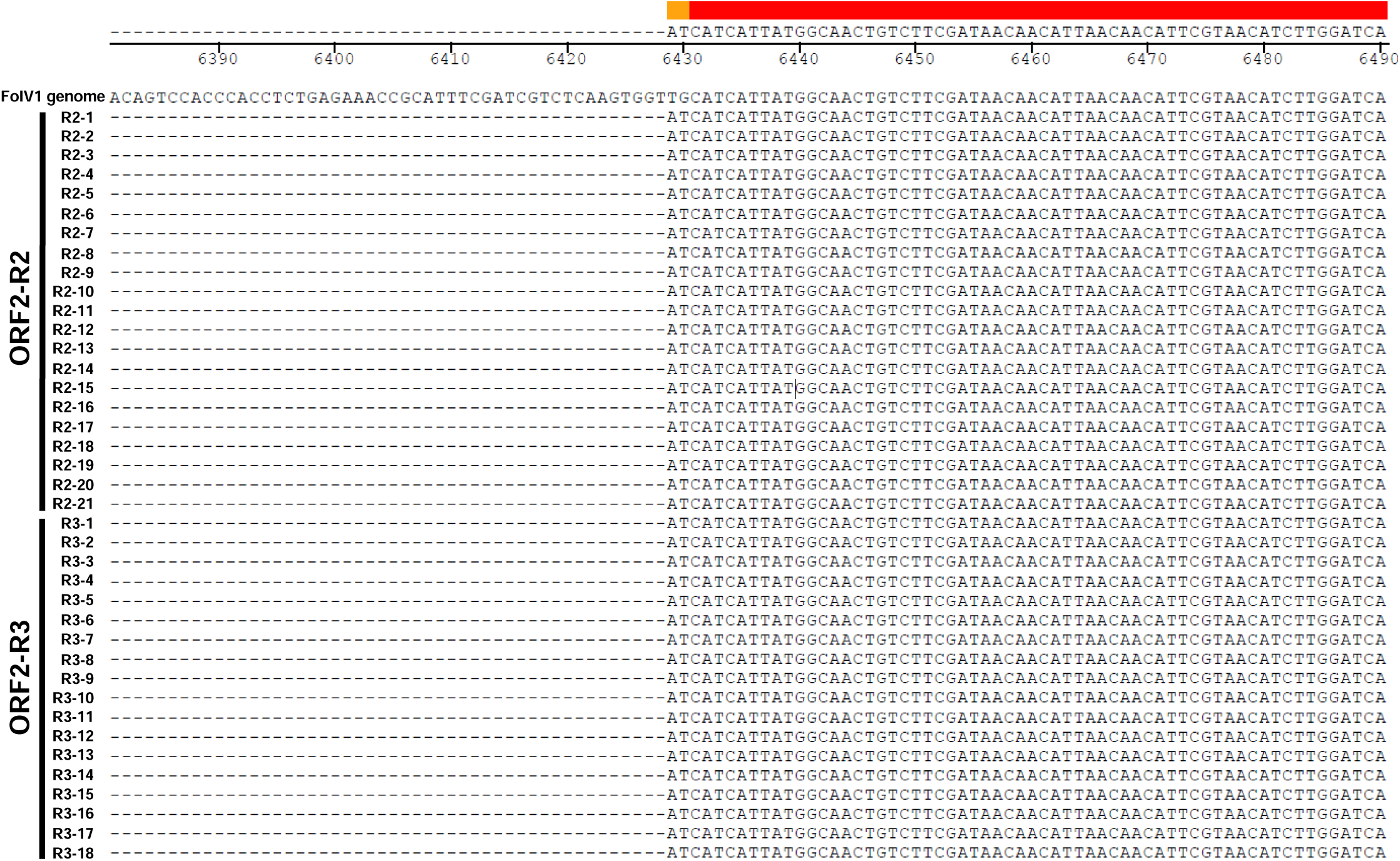
Alignment of the 5′-end sequences of 5′ RLM-RACE. The 39 obtained 5′-end sequences of 5′ RLM-RACE were sequenced and aligned using Megalign (DNASTAR).

**Supplementary Figure S4.**
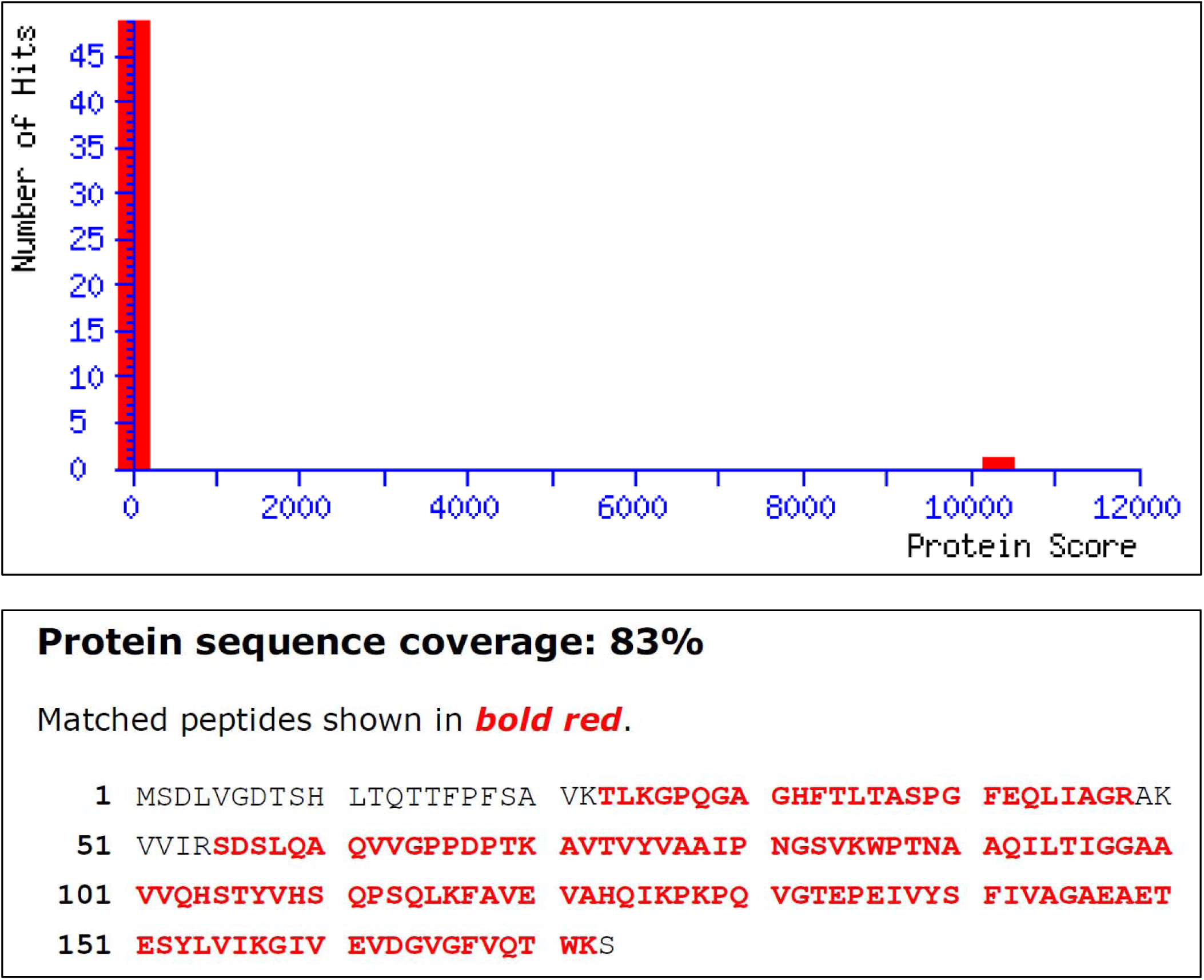
Mascot search result of 18 kDa protein of FoIV1 purification. Protein score: 10324 and matches 470 peptides of FoIV1 ORF4.

**Supplementary Figure S5.**
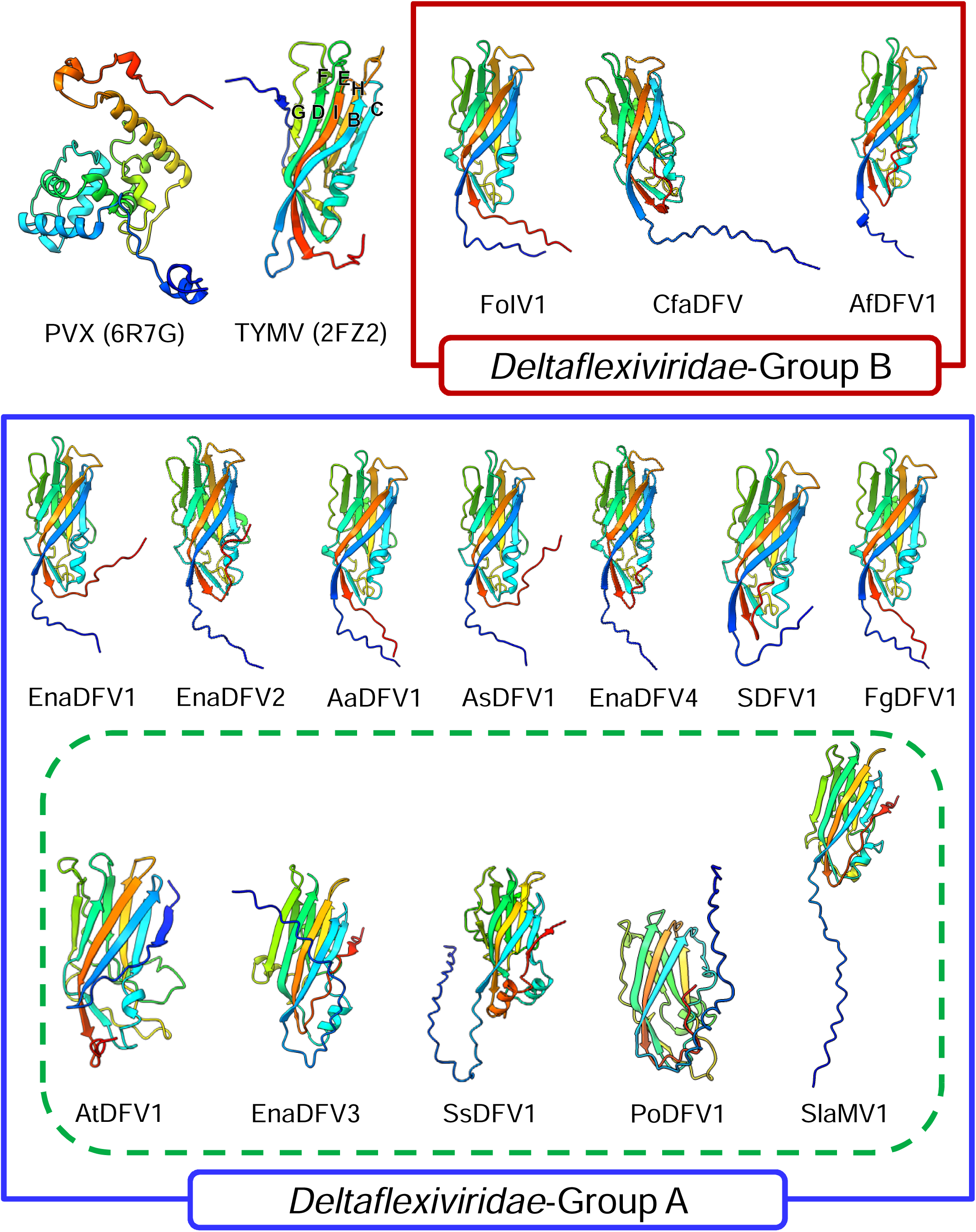
CP structures of the family *Deltaflexiviridae*-Group A & Group. **B.** PVX (6R7G) and TYMV (2FZ2) CP structures were obtained from the Protein Data Bank. CP structures of the family *Deltaflexiviridae*-Group A & Group B were predicted using AlphaFold2/ColabFold v1.5.5, then visualized UCSF ChimeraX v1.7.1.

**Supplementary Figure S6.**
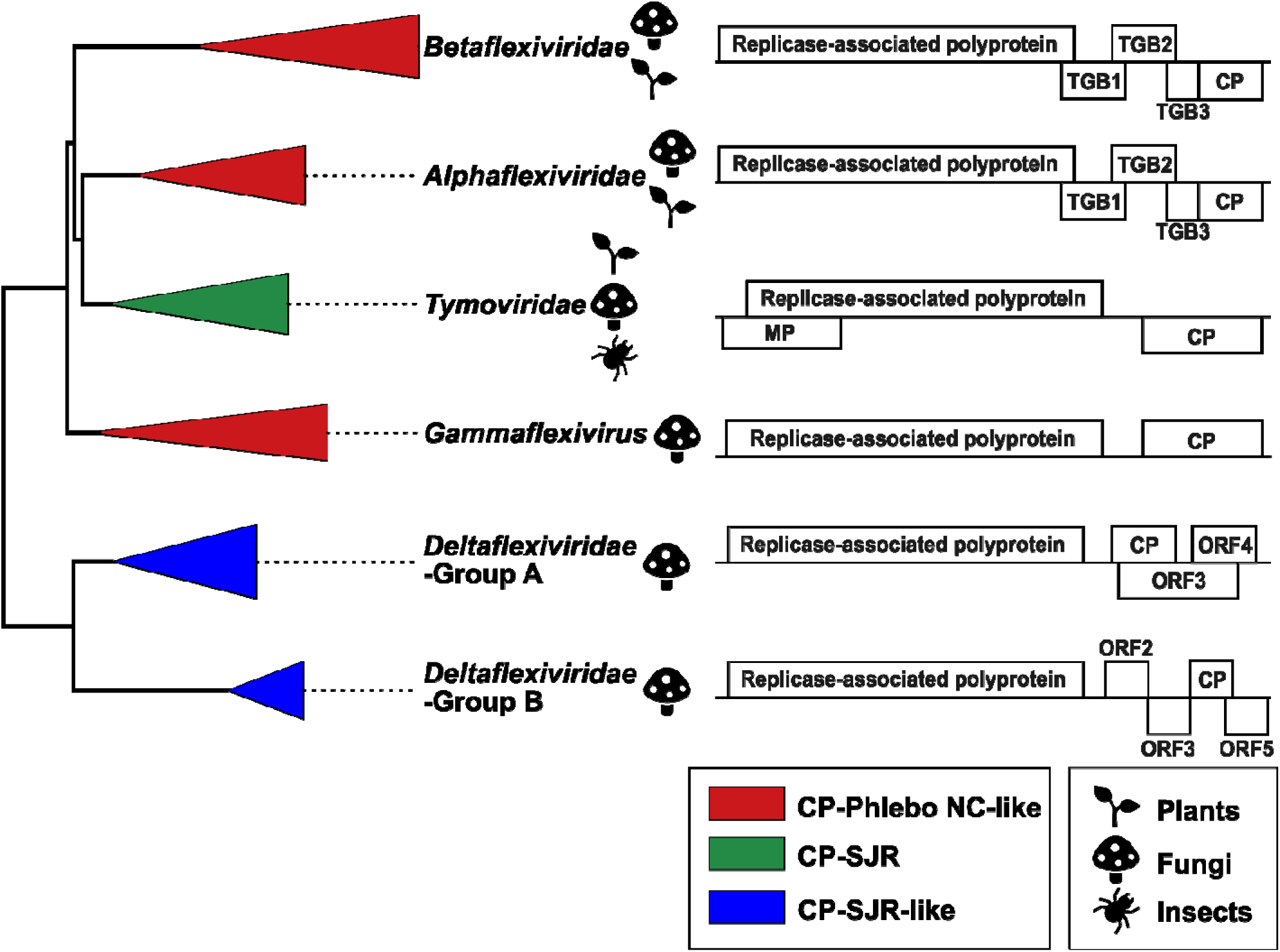
Evolution of *Tymovirales*. The dendrogram is based on the ML tree of Fig. 1C. Different families are shown as triangles colored according to the types of CPs: red, Phlebo NC-like CP; green, SJR CP; blue, SJR-like CP. Rough diagrams of representative virus genomes for each family showing encoded proteins. TGB, triple gene block; MP, movement protein; CP, coat protein; ORF, open reading frame of hypothetical protein.

**Supplementary Table S1.**
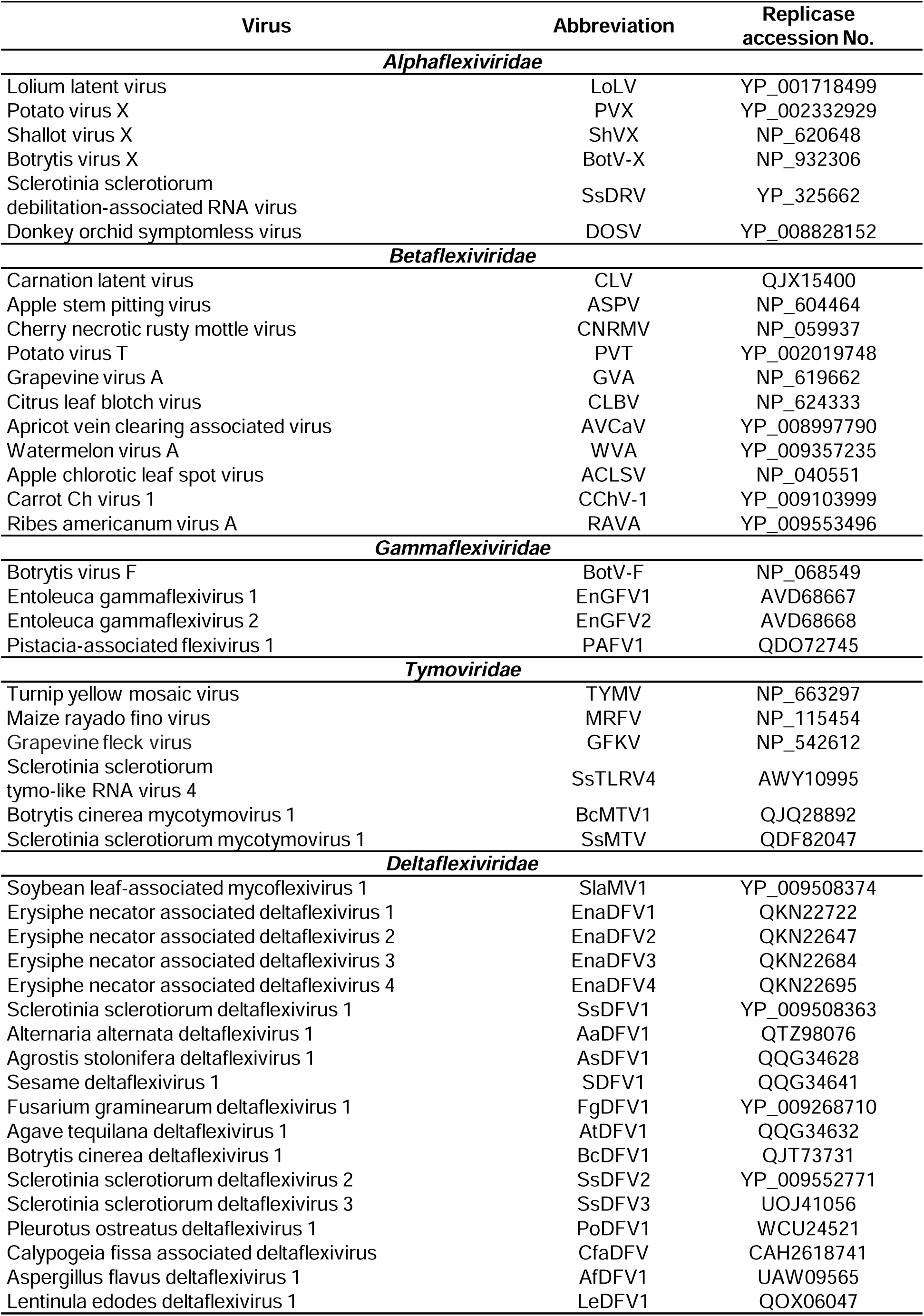
List of the amino acid sequences of replicases used in the phylogenetic analyses of this study.

**Supplementary Table S2.**
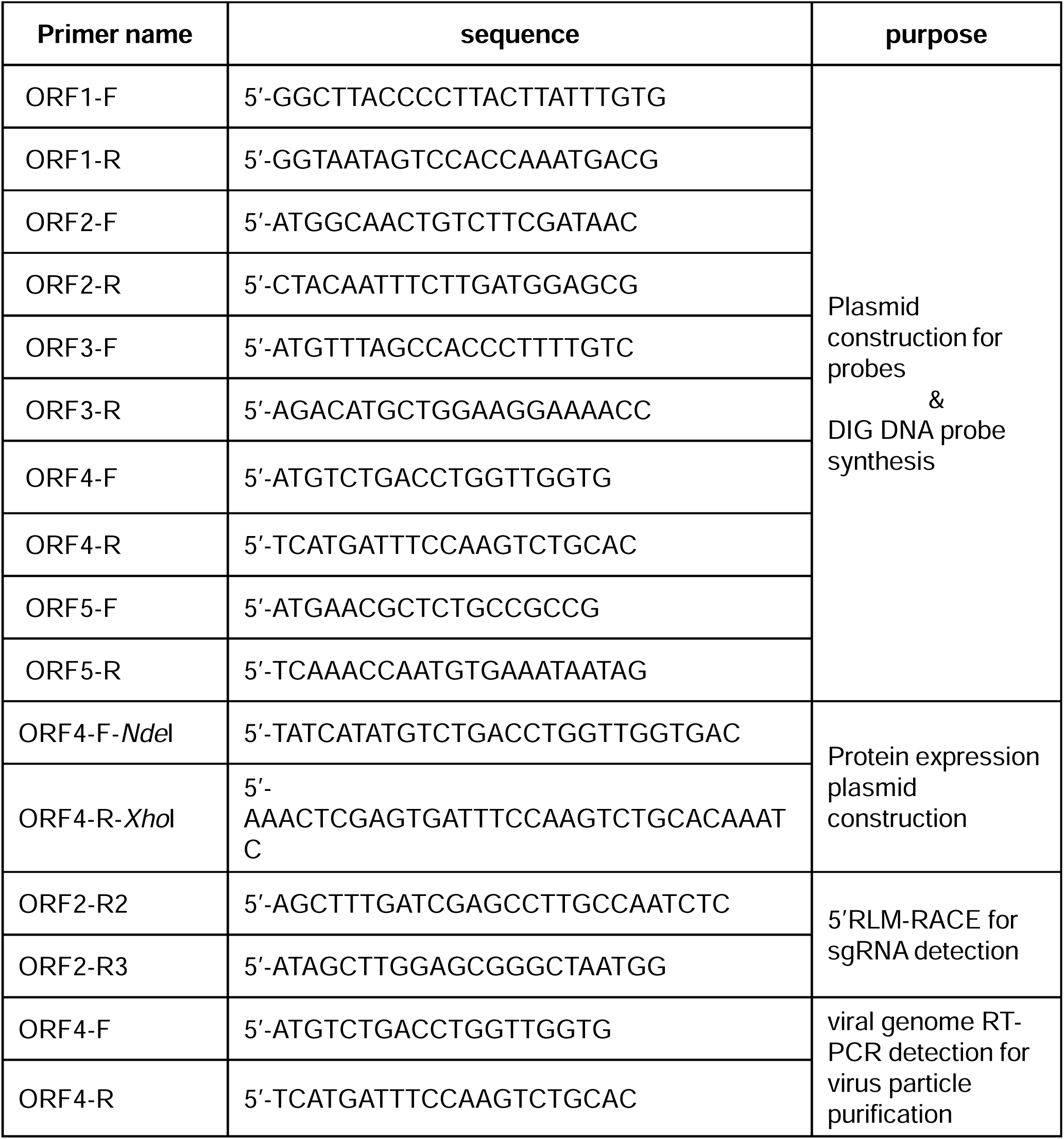
List of primers used in this study.

**Supplementary Table S3.**
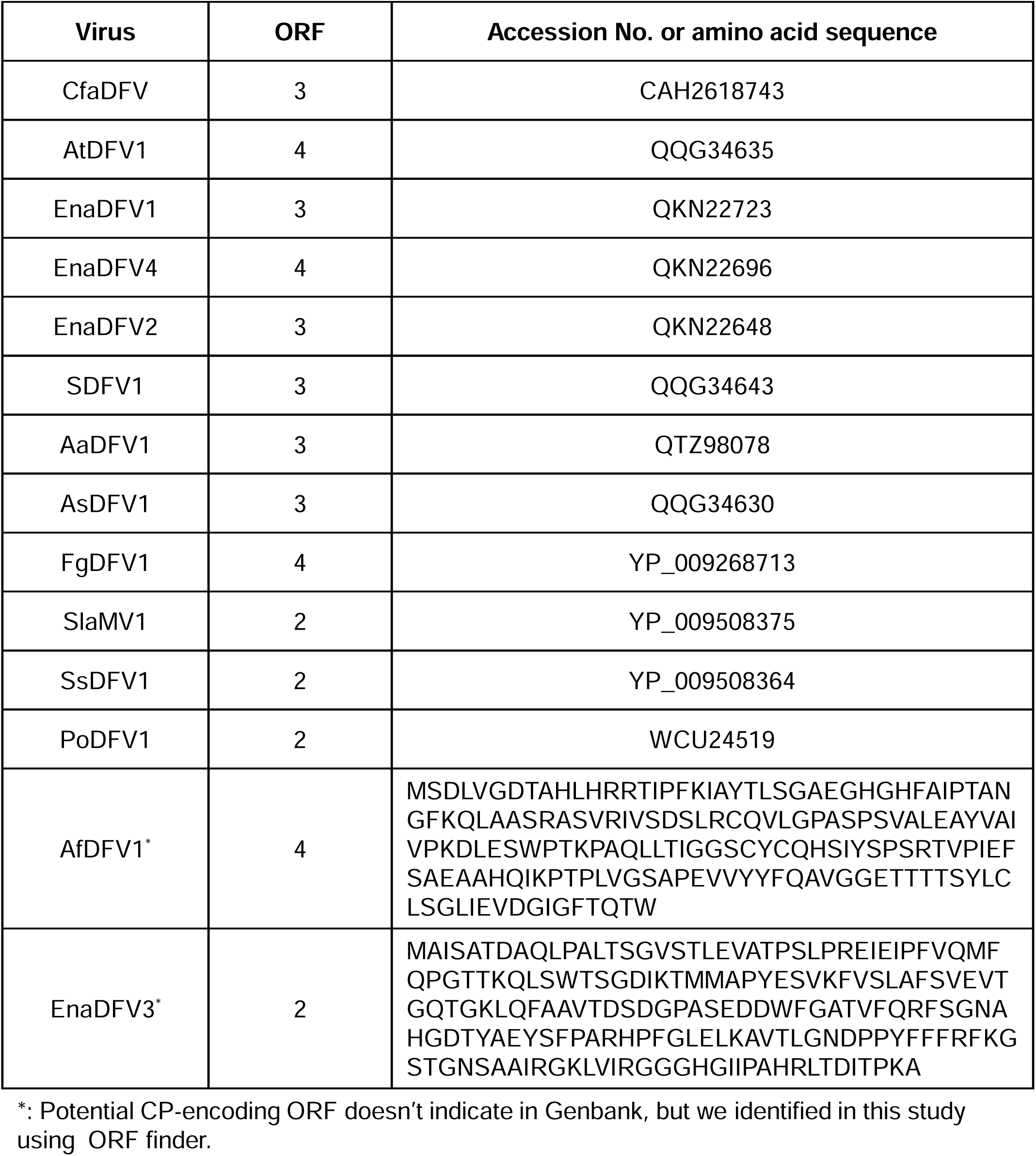
Predicted potential CP-encoding ORFs of deltaflexiviruses and FDFV2-related viruses.

